# Faster than thought: Detecting sub-second activation sequences with sequential fMRI pattern analysis

**DOI:** 10.1101/2020.02.15.950667

**Authors:** Lennart Wittkuhn, Nicolas W. Schuck

## Abstract

Neural computations are often anatomically localized and executed on sub-second time scales. Understanding the brain therefore requires methods that offer sufficient spatial and temporal resolution. This poses a particular challenge for the study of the human brain because non-invasive methods have either high temporal *or* spatial resolution, but not both. Here, we introduce a novel multivariate analysis method for conventional blood-oxygen-level dependent functional magnetic resonance imaging (BOLD fMRI) that allows to study sequentially activated neural patterns separated by less than 100 ms with anatomical precision. Human participants underwent fMRI and were presented with sequences of visual stimuli separated by 32 to 2048 ms. Probabilistic pattern classifiers were trained on fMRI data to detect the presence of image-specific activation patterns in early visual and ventral temporal cortex. The classifiers were then applied to data recorded during sequences of the same images presented at increasing speeds. Our results show that probabilistic classifier time courses allowed to detect neural representations and their order, even when images were separated by only 32 ms. Moreover, the frequency spectrum of the statistical sequentiality metric distinguished between sequence speeds on sub-second versus supra-second time scales. These results survived when data with high levels of noise and rare sequence events at unknown times were analyzed. Our method promises to lay the groundwork for novel investigations of fast neural computations in the human brain, such as hippocampal replay.

## Introduction

Many cognitive processes are underpinned by rapidly changing neural activation patterns. Most famously, memory and planning have been linked to fast replay of representation sequences in the hippocampus, happening approximately within 200 to 300 milliseconds (ms) while the animal is resting or sleeping [e.g., 1–9]. Similar events have been observed during behavior [10, 11], as well as outside of the hippocampus [12–17]. Likewise, internal deliberations during choice are reflected in alternations between orbitofrontal value representations that last less than 100 ms [18] and perceptual learning has been shown to result in sub-second anticipatory reactivation sequences in visual cortex [19–21]. Investigating fast-paced representational dynamics within specific brain areas therefore promises important insights into a variety of cognitive processes.

Such investigations are particularly difficult in humans, where signal detection must occur non-invasively, unless rare medical circumstances allow otherwise. How fast and anatomically localized neural dynamics can be studied using available neuroimaging techniques, in particular functional magnetic resonance imaging (fMRI), is therefore a major challenge for human neuroscience [for recent reviews, see e.g., 22, 23]. Here, we developed and experimentally validated a novel multivariate analysis method that allows to reveal the content and order of fast sequential neural events with anatomical specificity in humans using fMRI.

The main concern related to fMRI is that this technique measures neural activity indirectly through slow sampling of an extended and delayed blood-oxygen-level dependent (BOLD) response function [24–26] that can obscure temporal detail. Yet, the problems arising in BOLD fMRI might not be as insurmountable as they seem. First, BOLD signals from the same participant and brain region show reliable timing and last for several seconds. Miezin et al. [27], for instance, reported a between-session reliability of hemodynamic peak times in visual cortex of *r* ^2^ = .95 [see also 28, 29]. Even for closely timed events, the sequential order can therefore result in systematic differences in activation strength [30] that remain in the signal long after the fast sequence event is over, effectively mitigating the problems that arise from slow sampling. Second, some fast sequence events have properties that allow to detect them more easily. Replay events, in particular, involve reactivation of spatially tuned cells in the order of a previously travelled path. But these reactivated paths do not typically span the entire spatial environment and only involve a local subset of all possible places the animal could occupy [7, 8]. This locality means that even when measurement noise leads to partially re-ordered detection, or causes some elements of a fast sequence to remain undetected altogether, the set of detected representations will still reflect positions nearby in space. In this case, successive detection of elements nearby in space or time would still identify the fast process under investigation even under noisy conditions.

If fMRI analyses can fully capitalize on such effects, this could allow the investigation of fast sequential activations. One potential application of such methods would be hippocampal replay, a topic of intense recent interest [for reviews, see e.g., 23, 31–35]. To date, most replay research has studied the phenomenon in rodents because investigations in humans and other primates either required invasive recordings from the hippocampus [36–40], used techniques with reduced hippocampal sensitivity and spatial resolution [41–46], or investigated non-sequential fMRI activation patterns over seconds or minutes [47–51]. Recently, we have hypothesized that the properties of BOLD signals mentioned above should enable the investigation of rapid neural dynamics and identified fast sequential hippocampal pattern reactivation in resting humans using fMRI [52].

We extended this work in the present study by developing a modelling approach of multivariate fMRI pattern classification time courses and validating our method on experimentally controlled fast activation sequences in visual and ventral temporal cortex. As discussed above, we investigated the possibility to use fMRI to achieve (1) *order detection* and (2) *element detection* of fast activation sequences. The first effect, order detection, pertains to the presence of order structure in the signal that is caused by the sequential order of fast neural events. We evaluated this effect in two ways, first its impact on the relative strength of activations within a single measurement and second its consequences for the order across successive measurements. The second effect, element detection, quantifies to what extent fMRI allows to detect which elements were part of a sequence and which were not. While event detection is a standard problem in fMRI, we focused on the special case relevant to our question: detecting neural patterns of brief events that are affected by patterns from other sequence elements occurring only tens of milliseconds before or afterwards, causing backward and forward interference, respectively. Using full sequences of all possible elements in our experimental setup that tested sequence ordering, our design ensured that the two effects can be demonstrated independently, i.e., that the order effect could not have been a side effect of element detection. Our results demonstrate that fMRI with a conventional repetition time (TR) of 1.25 seconds (s) can be used to detect the elements and order of neural event sequences separated by only 32 ms. We also show that sequence detection can be achieved in the presence of high levels of signal noise and timing uncertainty, and is specific enough to differentiate fast sequences from activation patterns that could reflect slow conscious thinking.

## Results

To achieve full experimental control over fast activation patterns, we presented sequences of visual stimuli in a precisely timed and ordered manner. We then asked which aspects of the experimentally elicited fast neural processes are detectable from fMRI signals, and if detection is still possible when sequences occur embedded in noisy background activity at unknown times. We used multivariate pattern classifiers to analyze data from visual and ventral temporal cortex. Reflecting a common analytic scenario, classifiers were trained on fMRI data from individual events that proceeded at a slow pace (henceforth: *slow trials*, Fig. 1a) [cf. 42, 45, 50, 52]. We then applied the classifiers to (a) time points that contained sequences of events at different speeds (henceforth: *sequence trials*, Fig. 1b) and (b) trials involving varying numbers of event repetitions (henceforth: *repetition trials*, Fig. 1c), which allowed us to investigate sequence order and element detection, respectively. The analyses included *N* = 36 human participants who underwent two fMRI sessions each (four participants were excluded due to insufficient performance, see Methods and supplementary information (SI), Fig. S1a). Sessions were separated by 9 days on average (*SD* = 6 days, range: 1 – 24 days) and contained the trial types described below.

**Figure 1:**
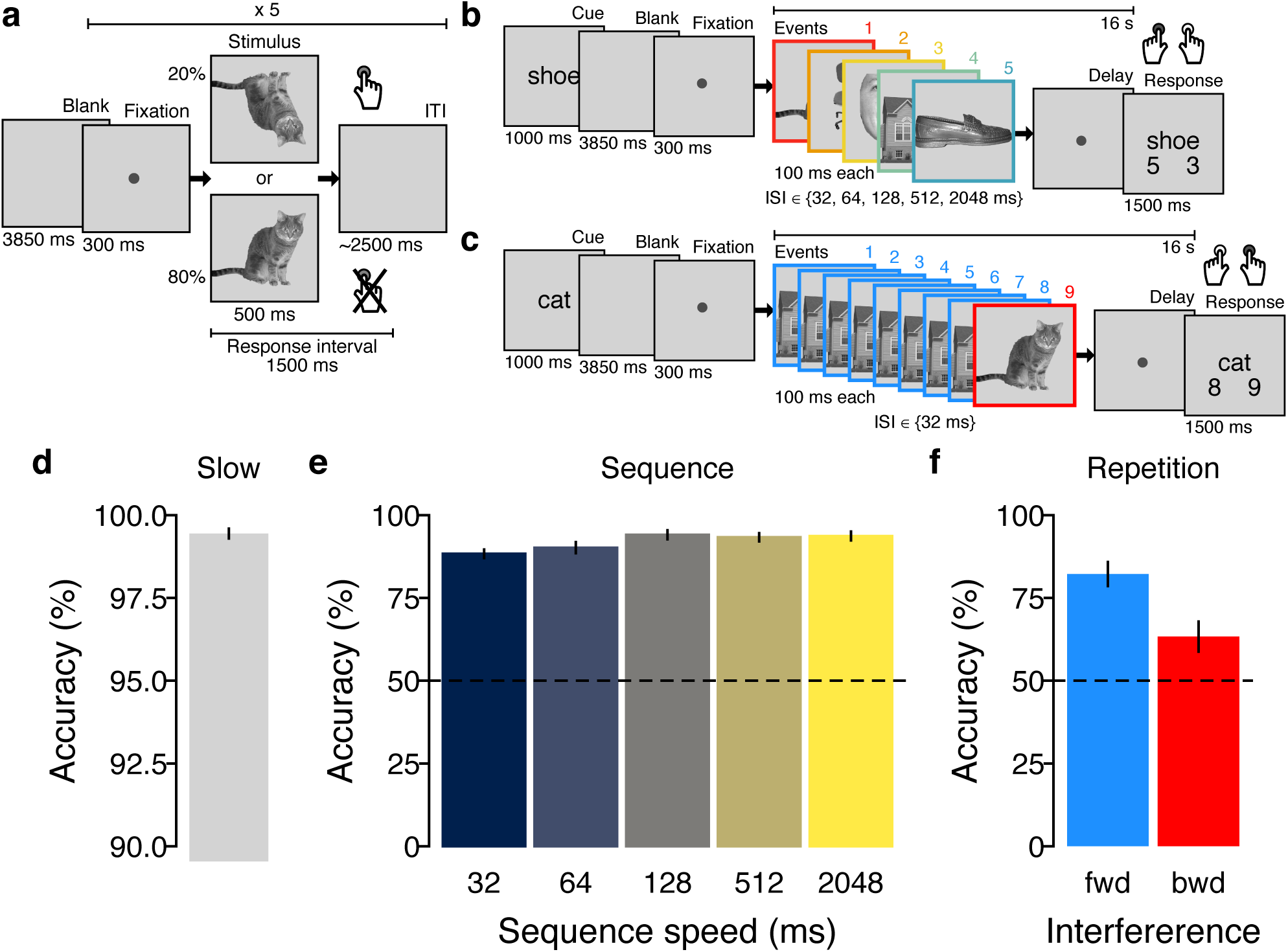
Task design and behavioral performance. **(a)** On slow trials, individual images were presented and inter-trial intervals (ITIs) were 2.5 s on average. Participants were instructed to detect upside-down visual stimuli (20% of trials) but not respond to upright pictures. Classifier training was performed on fMRI data from correct upright trials only. **(b)** Sequence trials contained five unique visual images, separated by five levels of inter-stimulus intervals (ISIs) between 32 and 2048 ms. **(c)** Repetition trials were always fast (32 ms ISI) and contained two visual images of which either the first or second was repeated eight times (causing backward and forward interference, respectively). In both task conditions, participants were asked to detect the serial position of a cued target stimulus in a sequence and select the correct answer after a delay period without visual input. One sequence or repetition trial came after five slow trials. **(d)** Mean behavioral accuracy (in %; y-axis) in upside-down slow trials. **(e)** Mean behavioral accuracy in sequence trials (in %; y-axis) as a function of sequence speed (ISI, in ms; x-axis). **(f)** Mean behavioral accuracy in repetition trials (in %; y-axis) as a function of which sequence item was repeated (fwd = forward, bwd = backward condition). All error bars represent ±1 standard error of the mean (SEM). The horizontal dashed lines in (e) and (f) indicate the 50% chance level.

### Training fMRI pattern classifiers on slow events

In slow trials, participants repeatedly viewed the same five images individually for 500 ms [images showed a cat, chair, face, house, and shoe; taken from 53]. Temporal delays between images were set to 2.5 s on average, as typical for task-based fMRI experiments [54]. To ensure that image ordering did not yield biased classifiers through biased pattern similarities [cf. 55], each possible order permutation of the five images was presented exactly once (120 sets of 5 images each). Participants were kept attentive by a cover task that required them to press a button whenever a picture was shown upside-down (20% of trials; mean accuracy: 99.44%; *t*_(35)_ = 263.27; *p* < .001, compared to chance; *d* = 43.88; Figs. 1d, S1a–c). Using data from correct upright slow trials, we trained five separate multinomial logistic regression classifiers, one for each image category [one-versus-rest; see Methods for details; cf. 53]. fMRI data were masked by a grey-matter-restricted region of interest (ROI) of occipito-temporal cortex, known to be related to visual object processing [11162 voxels in the masks on average; cf. 53, 56–58]. We accounted for hemodynamic lag by extracting fMRI data acquired 3.75 to 5 s after stimulus onset (corresponding to the fourth TR, see Methods). Cross-validated (leave-one-run-out) classification accuracy was on average 87.09% (*SD* = 3.50%; *p* < .001, compared to chance; *d* = 19.16; Fig. 2a). In order to examine the sensitivity of the classifiers to pattern activation time courses, we applied them to seven TRs following stimulus onset on each trial. This analysis confirmed delayed and distinct increases in the estimated probability of the true stimulus class given the data, peaking at the fourth TR after stimulus onset, as expected (Fig. 2b). The peak in probability for the true stimulus shown on the corresponding trial was significantly higher than the mean probability of all other stimuli at that time point (*t*s ≥ 17.89, *p*s < .001, *d*s ≥ 2.98; Bonferroni-corrected).

**Figure 2:**
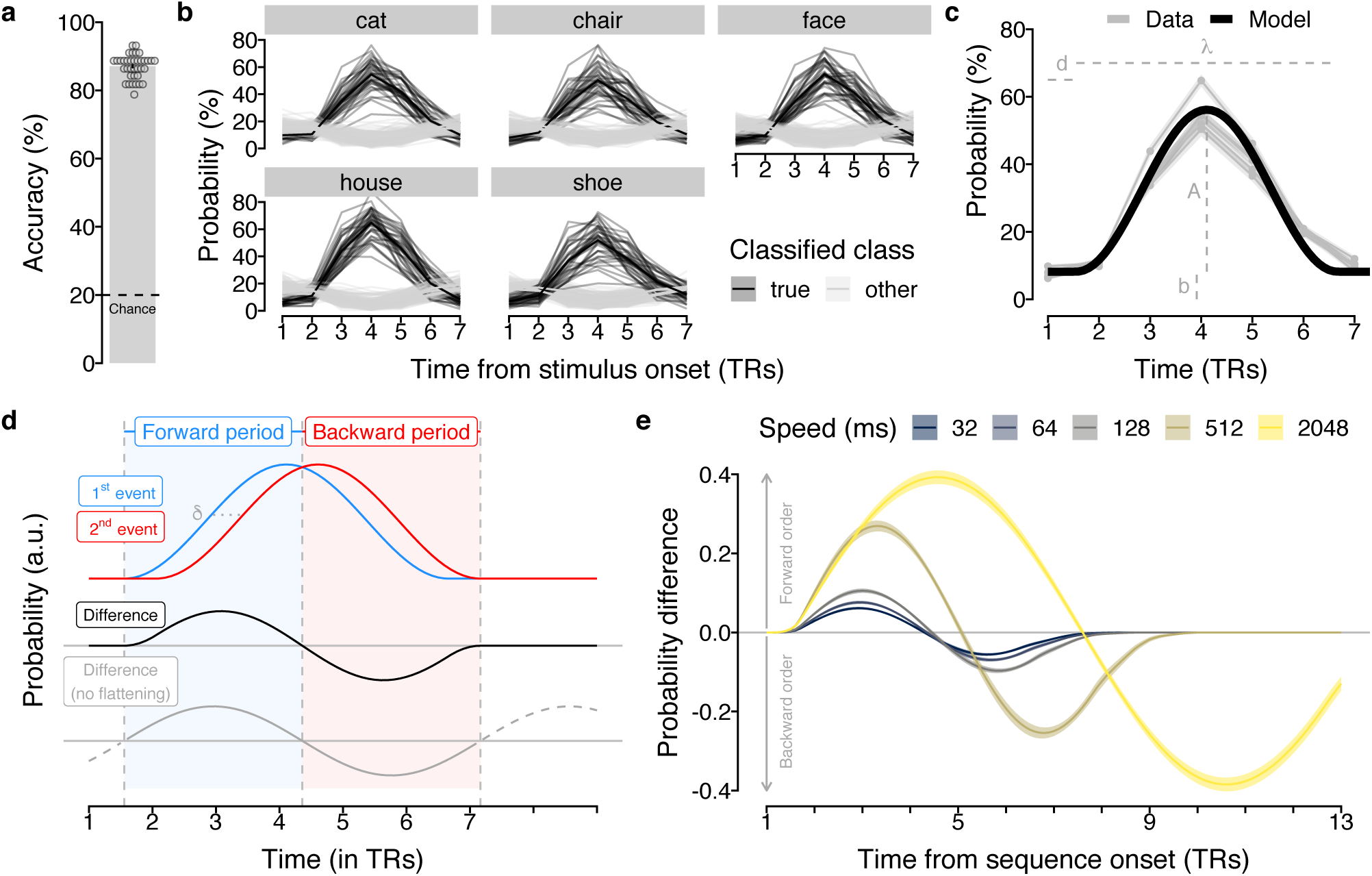
Classification accuracy and multivariate response functions. **(a)** Cross-validated classification accuracy in decoding the five unique visual objects in occipito-temporal data during task performance (in %; y-axis). Chance level is 20% (dashed line). Each dot corresponds to averaged data from one participant. Errorbar represents ±1 SEM. **(b)** Time courses (in TRs from stimulus onset; x-axis) of probabilistic classification evidence (in %; y-axis) for all five stimulus classes. Substantial delayed and extended probability increases for the stimulus presented (black lines) on a given trial (gray panels) were found. Each line represents one participant. **(c)** Average probabilistic classifier response for the five stimulus classes (gray lines) and fitted sine-wave response model using averaged parameters (black line). **(d)** Illustration of sinusoidal response functions following two neural events (blue and red lines) time-shifted by *δ* (dashed horizontal line). The resulting difference between event probabilities (black line) establishes a forward (blue area) and backward (red area) time period. The sine-wave approximation without flattened tails is shown in gray. **(e)** Probability differences between two time-shifted events predicted by the sinusoidal response functions depending on the event delays (*δ*) as they occurred in the five different sequence speed conditions (colors).

### Single event and event sequence modelling

The data shown in Fig. 2b highlight that multivariate decoding time courses are delayed and sustained, similar to single-voxel hemodynamics. We captured these dynamics elicited by single events by fitting a sine-based response function to the time courses on slow trials (a single sine wave flattened after one cycle, with parameters for amplitude *A*, response duration *λ*, onset delay *d* and baseline *b*, Figs. 2c, S2, see Methods). Based on this fit, we approximated expectations for signals during sequential events. The sequentiality analyses reported below essentially quantify how well successive activation patterns can be differentiated from one another depending on the speed of stimulus sequences. We therefore considered two time-shifted response functions and derived the magnitude and time course of *differences* between them. Based on the sinusoidal nature of the response function, the time course of this difference can be approximated by a single sine wave with duration *λ*_*δ*_ = *λ* + *δ*, where *δ* is the time between events and *λ* is the average fitted single event duration, here *λ* = 5.26 TRs (see Equations 4 and 5, Methods). This average parameter was used for all further analyses (Figs. 2c, 2d, see Methods). In this model, the amplitude is proportional to the time shift between events (until time shifts become larger than the time-to-peak of the response function). Consequently, after an onset delay (*d* = 0.56 TRs) the difference in probability of two time-shifted events is expected to be positive for the duration of half a cycle, i.e., 0.5*λ*_*δ*_ = 0.5(5.26 + *δ*) TRs, and negative for the same period thereafter. Three predictions arise from this model: (1) the first event will dominate the signal in earlier TRs and activation strengths will be proportional to the ordering of events during the sequential process; (2) in later TRs, the last sequence element will dominate the signal, and the activation strengths will be ordered in reverse; (3) the duration and strength of these two effects will depend on the fitted response duration and the timing of the stimuli as specified above (Fig. 2e, Equations 1–5, see Methods). For sequences with more than two items (like for sequence trials) *δ* is defined as the interval between the onsets of the first and last sequence item. We henceforth term the above mentioned early and late TRs the *forward* and *backward* periods, and consider all results below either separately for these phases, or for both relevant periods combined (calculating periods depending on the timings of image sequences and rounding TRs, see Methods).

### Detecting sequentiality in fMRI patterns following fast and slow neural event sequences

Our first major aim was to test detection of sequential order of fast neural events with fMRI. We therefore investigated above-mentioned *sequence trials* in which participants viewed a series of five unique images at different speeds (Fig. 1b). Sequence speed was manipulated by leaving either 32, 64, 128, 512 or 2048 ms between pictures, while images were always presented briefly (100 ms per image, total sequence duration 0.628–8.692 s). Sequences always contained each image exactly once. Every participant experienced 15 randomly selected image orders that ensured that each image appeared equally often at the first and last position of the sequence (all 120 possible orders counterbalanced across participants). The task required participants to indicate the serial position of a verbally cued image 16 s after the first image was presented. This delay between visual events and response allowed us to measure sequence-related fMRI signals without interference from following trials, while the upcoming question did not necessitate memorization of the sequence during the delay period. Performance was high even in the fastest sequence trials (32 ms: *M* = 88.33%, *SD* = 7.70, *p* < .001 compared to chance, *d* = 4.98), and only slightly reduced compared to the slowest condition (2048 ms: *M* = 93.70%, *SD* = 7.96, *p* < .001 compared to chance, *d* = 5.49, Figs. 1e,S1d).

We investigated whether sequence order detection was evident in the relative pattern activation strength within a single measurement. Examining the time courses of probabilistic classifier evidence during sequence trials (Fig. 3a) showed that the time delay between events was indeed reflected in sustained within-TR ordering of probabilities in all speed conditions. Specifically, immediately after sequence onset the first element (red line) had the highest probability and the last element (blue line) had the lowest probability. This pattern reversed afterwards, following the forward and backward dynamics that were predicted by the time-shifted response functions (Fig. 2d; forward and backward periods adjusted to sequence speed, see above and Methods). A TR-wise linear regression between the serial positions of the images and their probabilities confirmed this impression. In all speed conditions, the mean slope coefficients initially increased above zero (reflecting higher probabilities of earlier compared to later items) and decreased below zero afterwards (Figs. 3b, S4a). Considering mean regression coefficients during the predicted forward and backward periods, we found significant forward ordering in the forward period at ISIs of 128, 512 and 2048 ms (*t*s ≥ 2.83, *p*s ≤ .01, *d*s ≥ 0.47) and significant backward ordering in the backward period in all speed conditions (*t*s ≥ 3.94, *p*s < .001, *d*s ≥ 0.66, Fig. 3c). Notably, the observed time course of regression slopes on sequence trials (Fig. 3b) closely matched the time course predicted by our modeling approach (Fig. 2d), as indicated by strong correlations for all speed conditions between model predictions and the averaged time courses (Fig. 3d; Pearson’s *r*s ≥ .78, *p*s ≤ .001) as well as significant within participant correlations (Fig. 3e; Pearson’s *r*s ≥ .23, *t*s ≥ 3.67, *p*s ≤ .001 compared to zero, *d*s ≥ 0.61).

**Figure 3:**
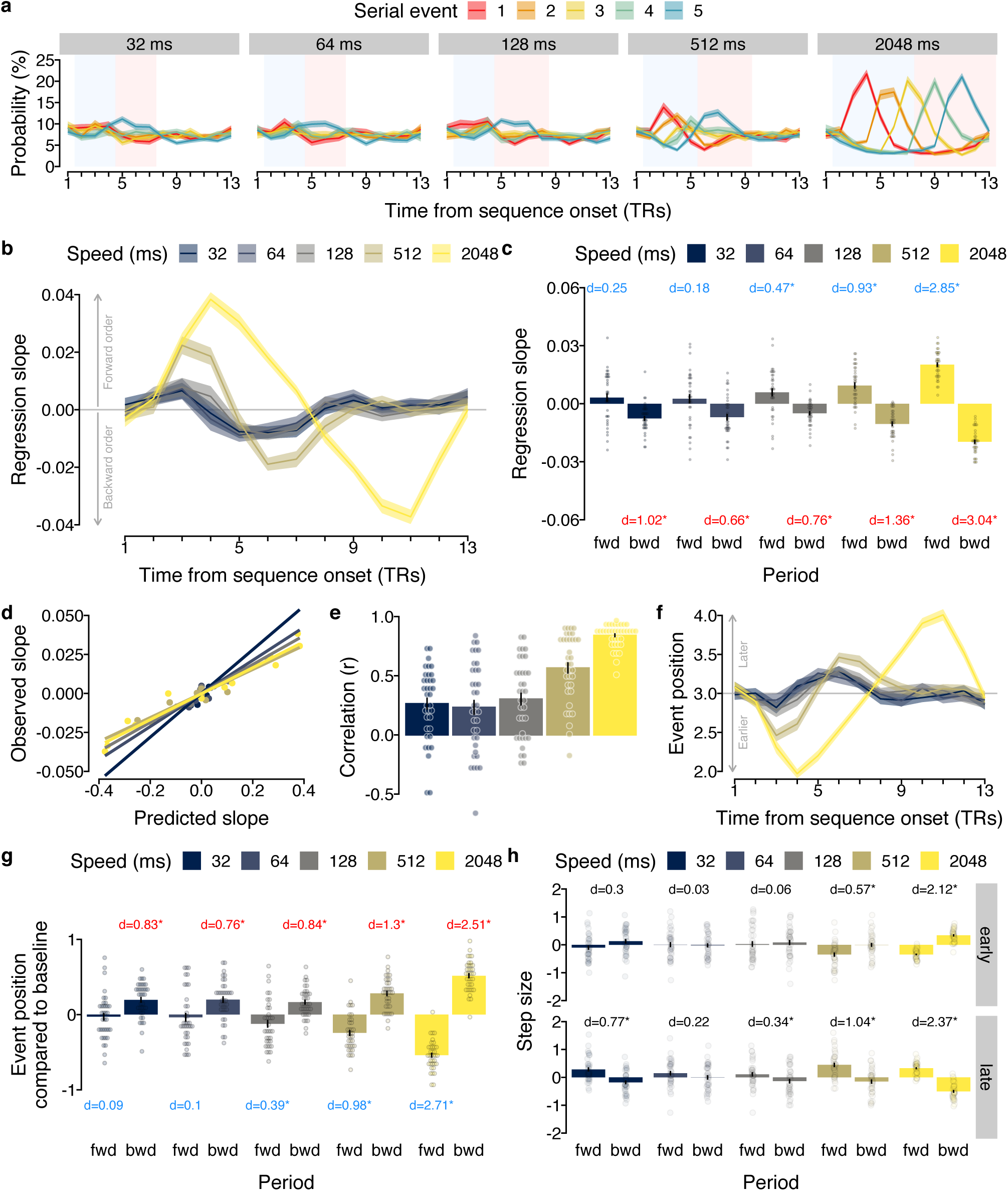
Sequence order is reflected in probability time courses. **(a)** Time courses (TRs from sequence onset) of classifier probabilities (%) per event (colors) and sequence speed (panels). Forward (blue) and backward (red) periods shaded as in Fig. 2d. **(b)** Time courses of mean regression slopes between event position and probability for each speed (colors). Positive / negative values indicate forward / backward sequentiality. **(c)** Mean slope coefficients for each speed (colors) and period (forward vs. backward; x-axis). Stars indicate significant differences from baseline. **(d)** Between-subject correlation between predicted (Fig. 2e) and observed (Fig. 3b) slopes. Each dot represents one TR. **(e)** Within-subject correlation between predicted and observed slopes as in (d). **(f)** Time courses of mean event position for each speed, as in (b). **(g)** Mean event position for each period and speed, as in (c) **(h)** Mean step sizes of early and late transitions for each period and speed. Stars indicate differences between periods, otherwise as in (c). Each dot represents data of one participant. Error bars/shaded areas represent ±1 SEM. Effect sizes indicated by Cohen’s *d*. Stars indicate *p* < .05, FDR-corrected. 1 TR = 1.25 s.

Choosing a different index of association like rank correlation coefficients (Figs. S3a–b, S4c) or the mean step size between probability-ordered events within TRs (Figs. S3c–d, S4d) produced qualitatively similar results (for details, see SI). Removing the sequence item with the highest probability at every TR also resulted in similar effects, with backward sequentiality remaining significant at all speeds (*p* ≤ .02) except the 128 ms condition (*p* = .10) and forward sequentiality still being evident at speeds of 512 and 2048 ms (*p* ≤ .002, Fig. S5a–b). To identify the drivers of the apparent asymmetry in detecting forward and backward sequentiality, we ran two additional control analyses and either removed the probability of the first or last sequence item (forward and backward periods adjusted accordingly). Removal of the first sequence item had little impact on sequentiality detection (Figs. S5c–d and SI), but removing the last sequence item markedly affected the results such that significant forward and backward sequentiality was only evident at speeds of 512 and 2048 ms (Figs. S5e–f and SI).

Next, we investigated evidence of pattern sequentiality across successive measurements, similar to Schuck and Niv [52]. Specifically, for each TR we only considered the decoded image with the highest probability and asked whether earlier images were decoded primarily in earlier TRs, and if later images were primarily decoded in later TRs. In line with this prediction, the average serial position fluctuated in a similar manner as the regression coefficients, with a tendency of early positions to be decoded in early TRs, and later positions in later TRs (Fig. 3f). The average serial position of the decoded images was therefore significantly different between the predicted forward and backward period at all sequence speeds (all *p*s < .001, Figs. 3g, S4d). Compared to baseline (mean serial position of 3), the average serial position during the forward period was significantly lower for speeds of 128, 512 and 2048 ms (all *p*s ≤ .03). The average decoded serial position at later time points was significantly higher compared to baseline in all speed conditions, including the 32 ms condition (all *p*s < .001). Thus, earlier images were decoded earlier after sequence onset and later images later, as expected. This sequential progression through the involved sequence elements had implications for transitions between consecutively decoded events. Initially, when early elements begin to dominate the signal in the first half of the forward period (henceforth *early*), the position of decoded sequence items decreased relative to baseline. During the first half of the backward period, however, the decoded serial positions increased, reflecting the ongoing progression through all sequence elements from first to last. The reverse was true during the second half of both periods (henceforth *late*): positions began to increase in the forward period, but during the second half of the backward period, the decoded positions were about to return back to baseline from the last decoded item, thus decreasing again. To verify this effect, we computed the step sizes between consecutively decoded serial events as in Schuck and Niv [52]. For example, observing a 2→4 transition of decoded events in consecutive TRs would correspond to a step size of +2, while a 3→2 transition would reflect a step size of –1. In line with the above-mentioned predictions, the step sizes of *early* transitions were significantly more forward directed in the forward as compared to the backward period for speed conditions of 512 and 2048 ms (*p*s ≤ .005, Fig. 3h). Average step sizes of *late* transitions, in contrast, were negative directed in the forward period and vice versa in the backward period, differing in all speed conditions (*p*s ≤ .05, Fig. 3h), except the 64 ms condition (*p* = .19). This analysis suggests that transitions between decoded items reflect the gradual progression through all sequence events, even when events were separated only by tens of milliseconds.

### Detecting sequence elements: asymmetries and interference effects

We next turned to our second main question, asking whether we can detect which patterns were part of a fast sequence and which were not. To this end, we investigated classification time courses in repetition trials, in which only two out of the five possible images were shown. Crucially, one image was repeated, while the other one was shown only once. Embedding one briefly displayed image into the context of a repeated image allowed us to study to what extent another activation can interfere with the detection of a brief activation pattern of interest. Repeating the interfering image eight times allowed us to study this phenomenon in a worst case scenario by exaggerating the interference effect. Finally, varying whether the second or first item is short allowed us to investigate if the ability to detect sequence elements is asymmetrical, and possibly favors the detection of late over early events. Specifically, if the first image was shown briefly once and followed immediately by eight repetitions of a second image, the dominant second image will interfere with the detection of the first image (henceforth *forward interference* condition, since the forward phase suffers from interference). If, on the other hand, the first image was repeated eight times and the second image was shown once, the first image will be dominant and possibly interfere with the backwards phase (henceforth *backward interference* condition). Comparing the forward and backward conditions therefore allowed closer assessment of asymmetries, which had become apparent in the results presented above (Fig. 3).

In all cases, images were separated by only 32 ms. As before, we applied the classifiers trained on slow trials to the data acquired in repetition trials, to obtain the estimated probability of every class given the data for each TR (Figs. 4a, S7). The expected relevant time period was determined to be from TRs 2 to 7 and used in all analyses (see rectangular areas in Fig. 4a). Participants were kept attentive by the same cover task used in sequence trials (Fig. 1c). Average behavioral accuracy was high on repetition trials (*M* = 73.46%, *SD* = 9.71%; Figs. 1f, S1a) and clearly differed from a 50% chance-level (*t*_(35)_ = 14.50, *p* < .001, *d* = 2.42). Splitting up performance into forward and backward interference trials showed performance above chance level in both conditions (*M* = 82.22% and *M* = 63.33%, respectively, *p*s ≤ .003, *d*s ≥ 0.49, Fig. 1f). Additional conditions with intermediate levels of repetitions are reported in the SI (Fig. S1e).

**Figure 4:**
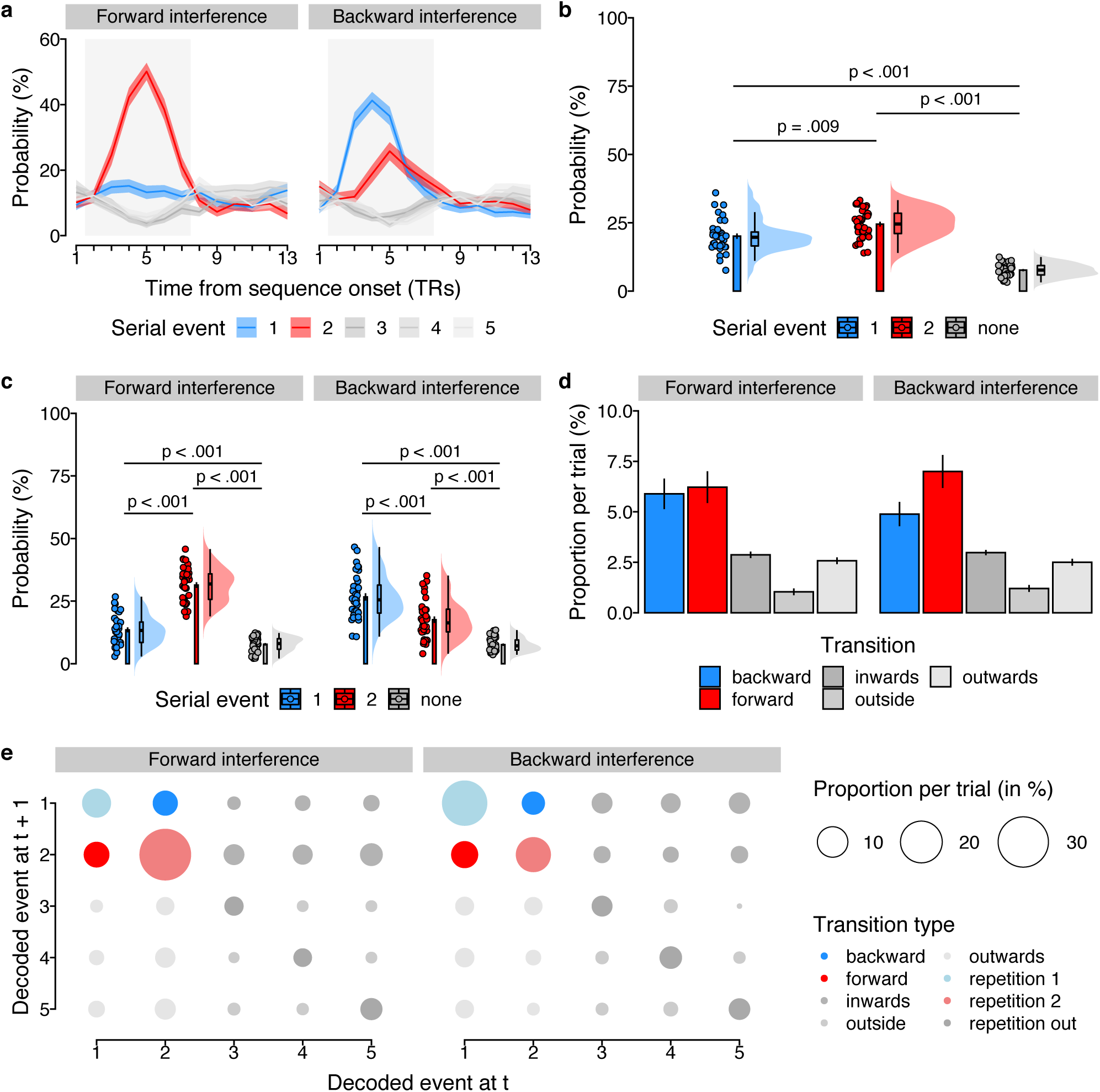
Ordering of two-item pairs on repetition trials. **(a)** Time courses (in TRs from sequence onset; x-axis) of probabilistic classifier evidence (in %) in repetition trials, color-coded by event type (first/second/non-sequence, see legend). Data shown separately for forward (left) and backward (right) interference conditions. Gray background indicates relevant time period independently inferred from response functions (Fig 2d). Shaded areas represent ±1 SEM. 1 TR = 1.25 s. **(b)** Mean probability of event types averaged across all TRs in the relevant time period, as in (a). Each dot represents one participant, the probability density of the data is shown as rain cloud plots [cf. 59]. Boxplots indicate the median and interquartile range. The barplots show the sample mean and errorbars indicate ±1 SEM. **(c)** Average probability of event types, separately for conditions as in (a), plots as in (b). **(d)** Mean trial-wise proportion of each transition type, separately for forward/backward conditions, as in (a). **(e)** Transition matrix of decoded images indicating mean proportions per trial, separately for the forward and backward condition (left/right). Transition types highlighted in colors (see legend).

We first asked whether our classifiers indicated that the two events that were part of the sequence were more likely than items that were not part of the sequence. Indeed, the event types (first, second, non-sequence) had significantly different mean decoding probabilities, with sequence items having a higher probability (first: *M* = 20.09%; second: *M* = 24.52%) compared to non-sequence items (*M* = 7.68%; both *ps* < .001, corrected; main effect: *F*_2,53.51_ = 106.94, *p* < .001, Fig. 4b). Moreover, the probability of decoding within-sequence items depended on their position as well as the their duration (number of repetitions). Considering both interference conditions revealed a main effect of event type, *F*_2,40.18_ = 135.88, *p* < .001, as well as an interaction between event type and duration, *F*_2,105.0_ = 123.35, *p* < .001, but no main effect of duration, *p* = .70 (Fig. 4c). This indicated that the forward phase suffered from much stronger interference than the backwards phase. In the *forward interference* condition the longer second event had an approximately 18% higher probability than the first event (31.44% vs 13.52%, *p* < .001), whereas in the *backward interference* condition the first event had an only 9% higher probability than the second (26.67% vs. 17.60%, *p* < .001, corrected). Thus, item detection is impacted more by succeeding than preceding activation patterns, leading to the increased dominance of the last item in sequence trials particularly in the fast conditions (Fig. 3a). Importantly, however, both sequence elements still differed from non-sequence items even under conditions of interference (forward: 7.76% and backward: 7.59%, respectively, all *p*s < .001, corrected), indicating that sequence element detection remains possible under such circumstances. Using data from all TRs revealed qualitatively similar significant effects (*p* < .05 for all but one test after correction, see SI). Repeating all analyses using proportions of decoded classes (the class with the maximum probability was considered decoded at every TR), or considering all repetition trial conditions, also revealed qualitatively similar results). Thus, brief events can be detected despite significant interference.

We next asked which implications these findings have for the observed pattern transitions [cf. 52]. To this end, we analyzed the trial-wise proportions of transitions between consecutively de-coded events, and asked whether forward transitions between sequence items were more likely than transitions between a sequence and a non-sequence item (outward transitions) or between two non-sequence items (outside transition; details see Methods). This analysis revealed that forward transitions (6.22%) were more frequent than both outward transitions (2.57%), and outside transitions (1.04%, both *p*s < .001, corrected; Fig. 4d) in the *forward interference* condition. The same was true in the *backward interference* condition (forward transitions: 7.00%; outward transitions: 2.50%; outside transitions: 1.20%, all *p*s < .001). The full transition matrix is shown in Fig. 4e.

Together, the results from repetition trials indicated that (1) within-sequence items could be clearly detected despite interference from other sequence items, (2) event detection was asymmetric, such that items occurring at the end of sequences can be detected more easily than those occurring at the beginning and (3) sequence item detection leads to within sequence pattern transitions.

### Detecting sparse sequence events with lower signal-to-noise ratio (SNR)

The results above indicate that detection of fast sequences is possible if they are under experimental control. In most applications of our method, however, this will not be the case. When detecting replay, for instance, sequential events will occur spontaneously during a period of noise. We therefore next assessed the usefulness of our method under such circumstances.

We first characterized the behavior of sequence detection metrics during periods of noise. To this end, we applied the logistic regression classifiers to fMRI data acquired from the same participants (*N* = 32 out of 36) during a 5-minute (233 TRs) resting period before any task exposure in the scanner. Classifier probabilities during rest fluctuated wildly, often with a single category having a high probability, while all other categories had probabilities close to zero. During fast sequence periods, in contrast, the near-simultaneous activation of stimulus-driven activity led to reduced probabilities, such that category probabilities tended to be closer together and less extreme. In consequence, the average standard deviation of the probabilities per TR during rest and slow (2048 ms) sequence periods was higher (*M* = 0.23 and *M* = 0.22, respectively) compared to the average standard deviation in the fast sequence condition (32 ms; *M* = 0.20; *t*s ≥ 4.02; *p*s ≤ .001; *d*s ≥ 0.71; Fig. 5a).

**Figure 5:**
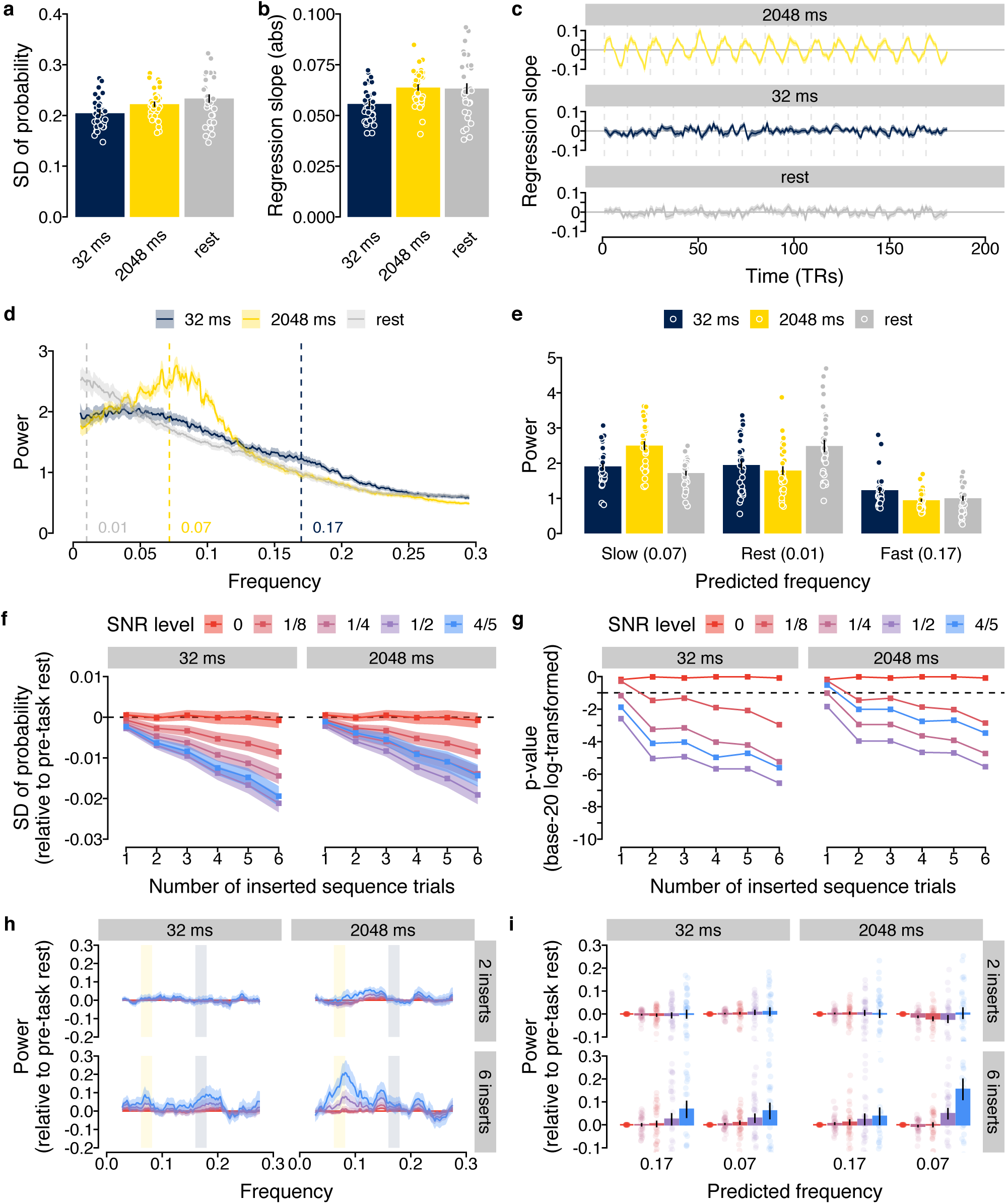
Detecting sparse sequence events with lower SNR. **(a)** Mean standard deviation of classifier probabilities in rest and sequence data. **(b)** Mean absolute regression slopes, as in (a). **(c)** Time courses of regression slopes in rest and sequence data. Vertical lines indicate trial boundaries. **(d)** Normalized frequency spectra of regression slopes in rest and sequence data. Annotations indicate predicted frequencies based on Eqn. 5. **(e)** Mean power of predicted frequencies in rest and sequence data, as in (a). Each dot represents data from one participant. **(f)** Mean standard deviation of rest data including a varying number of SNR-adjusted sequence events (fast or slow). Dashed line indicates indifference from sequence-free rest. **(g)** Base-20 log-transformed *p*-values of *t*-tests comparing the standard deviation of probabilities in (f) with sequence-free rest. Dashed line indicates *p* = .05. **(h)** Frequency spectra of regression slopes in SNR-adjusted sequence-containing rest relative to sequence-free rest. Rectangles indicate predicted frequencies, as in (d). **(i)** Mean relative power of predicted frequencies in SNR-adjusted sequence-containing rest. All *p*s FDR-corrected. Shaded areas / error bars represent ± 1 SEM. 1 TR = 1.25 s.

As before, we next fitted regression coefficients through the classifier probabilities of the rest data and, for comparison, to concatenated data from the 32 ms and 2048 ms sequence trials (Fig. 5b–c). As predicted by our modelling approach (Fig. 2e), and shown in the previous section (Fig. 3b), the time courses of regression coefficients in the sequence conditions were characterized by rhythmic fluctuations whose frequency and amplitude differed between speed conditions (Fig. 5c). To quantify the magnitude of this effect, we calculated frequency spectra of the time courses of the regression coefficients in rest and concatenated sequence data (Fig. 5d; using the Lomb-Scargle method [e.g., 60] to account for potential artefacts due to data concatenation, see Methods). This analysis revealed that frequency spectra of the sequence data differed from rest frequency spectra in a manner that depended on the speed condition (Fig. 5d–e). As foreshadowed by our model, power differences appeared most pronounced in the predicted frequency ranges (Fig. 5e; *p*s ≤ .02; see Eqn. 5 and Methods).

Finally, we asked whether these differences would persist if (a) only few sequence events occurred during a 5-minute rest period, while (b) their onset was unknown and (c) their SNR was lower. To this end, we synthetically generated data containing a variable number of sequence events that were inserted at random times into the resting state data acquired before any task exposure. Specifically, we inserted between 1 and 6 sequence events into the rest period by blending rest data with TRs recorded in fast (32 ms) or slow (2048 ms) sequence trials (12 TRs per trial, random selection of sequence trials and insertion of time points, without replacement). To account for possible SNR reductions, the inserted probability time courses were multiplied by a factor *κ* of 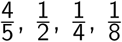 or 0 and added to the probability time courses of the inversely scaled (1 – *κ*) resting state data. Effectively, this led to a step-wise reduction of the inserted sequence signal from 80% to 0%, relative to the SNR obtained in the experimental conditions reported above.

As expected, differences in above-mentioned standard deviation of the probability gradually increased with both the SNR level and the number of inserted sequence events when either fast or slow sequences were inserted (Fig. 5f). In our case this led significant differences to emerge with one insert and an SNR reduced to 12.5% in both the fast and slow condition (Fig. 5g; comparing against 0, the expectation of no difference with a conventional false positive rate *α* of 5%; all *p*s false discovery rate (FDR)-adjusted).

Importantly, the presence of sequence events was also reflected in the frequency spectrum of the regression coefficients. Inserting fast event sequences into rest led to power increases in the frequency range indicative of 32 ms events (∼ 0.17 Hz, Fig. 5f, left panel), in line with our findings above. This effect again got stronger with higher SNR levels and more sequence events. Inserting slow (2048 ms) sequence events into the rest period showed a markedly different frequency spectrum, with an increase around the frequency predicted for this speed (∼ 0.07 Hz, Fig 5f, right panel). Comparing the power around the predicted frequency (±0.01 Hz) of both speed conditions indicated significant increases in power compared to sequence-free rest when six sequence events were inserted and the SNR was reduced to 80% (*t*s ≥ 2.11, *p*s ≤ .04, *d*s ≥ 0.37). Hence, the presence of spontaneously occurring sub-second sequences during rest can be detected in the frequency spectrum of our sequentiality measure, and distinguished from slower second-scale sequences that might reflect conscious thinking.

## Discussion

We demonstrated that BOLD fMRI can be used to localize sub-second neural events sequences non-invasively in humans. We combined probabilistic multivariate pattern analysis with time course modelling and investigated human brain activity recorded following the presentation of sequences of visual objects at varying speeds. In the fastest case a sequence of five images was displayed within 628 ms (32 ms between pictures). Even when using a TR of only 1.25 s (achievable with conventional multi-band echo-planar imaging), the image order could be detected from activity patterns in visual and ventral temporal cortex. Detection of briefly presented sequence items was also possible when their activation was affected by interfering signals from a preceding or subsequent sequence item and could be differentiated from images that were not part of the sequence. Our results withstood several robustness tests, but also indicated that detection is biased to most strongly reflect the last event of a sequence. Analyses of augmented resting data, in which neural event sequences occurred rarely, at unknown times, and with reduced signal strength, showed that our method could detect sub-second sequences even under such adverse conditions. Moreover, we showed that frequency spectrum analyses allow to distinguish sub-second from supra-second sequences under such circumstances. Our approach therefore promises to expand the scope of BOLD fMRI to fast, sequential neural representations by extending multivariate decoding approaches into the temporal domain, in line with our previous findings [52].

One important potential application of our method is the study of replay, the temporally compressed sequential reactivation of neural representations in hippocampal and neocortical areas that subserves memory consolidation, planning, and decision-making [for reviews, see e.g., 31, 33, 61, 62]. Previous fMRI studies in humans [for reviews see e.g., 23, 63] measured non-sequential reactivation as increased similarity of multivoxel patterns during experience and extended post-encoding rest compared to pre-encoding baseline [47–49, 51, 64–68] or functional connectivity of hippocampal, cortical and dopaminergic brain structures that support post-encoding systems-level memory consolidation [65–67, 69–71]. In the current study we open the path to extend this fMRI research towards an understanding of the speed and sequential nature of the observed phenomena.

Our fMRI-based approach has advantages as well as disadvantages compared to existing electroencephalography (EEG) and magnetoencephalography (MEG) approaches [42, 44, 45]. In particular, it seems likely that our method has limited resolution of sequence speed. While we could distinguish between supra- and sub-second sequences, a finer distinction was not feasible. Yet, EEG and MEG investigations suggest that the extent of temporal compression of previous experience is an important aspect of replay and other reactivation phenomena [43, 72–75]. In addition, the differential sensitivity to activity depending on sequence position complicates interpretations of findings, and can lead to statistical aliasing of sequences with the same start and end elements but different elements in the middle. Finally, because a single sequence causes forward and backward ordering of signals, it can be difficult to determine the direction of a hypothesized sequence. The major advantage of fMRI is that it does not suffer from the low sensitivity to hippocampal activity and limited ability to anatomically localize effects that characterizes EEG and MEG. This is particularly important in the case of replay, which is hippocampus-centered but co-occurs with fast sequences in other parts of the brain including primary visual cortex [12], auditory cortex [15], prefrontal cortex (PFC) [13, 14, 16, 17, 76], entorhinal cortex [77–79], and ventral striatum [80]. Importantly, replay events occurring in different brain areas might not be mere copies of each other, but can differ regarding their timing, content and relevance for cognition [e.g., 16, 17]. Precise characterization of replay events occurring in different anatomical regions is therefore paramount. Because EEG and MEG cannot untangle the co-occurring events and animal research is often restricted to a single recording site, much remains to be understood about the distributed and coordinated nature of replay.

Finally, our study provides insights for future research. First, the bias towards later sequence events has to be taken into account when analyzing data for which the ground truth is not known. Second, we have shown that the mere fact that detecting which elements where part of a sequence is beneficial if sequences mostly contain a local subset of all possible events. Thus, experimental setups with a larger number of possible events will be useful. At the same time, a larger number of to be decoded events will likely impair baseline classification accuracy, which in turn impairs sequence detection. Researchers should thus take the trade-off between these two aspects into account. Third, several other factors emerged that could influence the success of future investigation: the sampling rate (the TR), the choice of brain region and the properties of the resulting hemodynamic response functions (HRFs) [22]. It should be noted, however, that an increased sampling rate will only partially increase power, since the extended HRF duration ensures measurement opportunities up to 10 s after the sequence. Moreover, the choice of brain region will impact results only if the stability of the HRF within that brain region is low, whereas between-region differences between HRF parameters might have less impact. But HRF stability is generally high [29, 81–83], and previous research noting this fact has therefore already indicated possibilities of disentangling temporally close events [27–30, 84, 85]. Our approach has shown how using multivariate and modelling approaches can help exploit these HRF properties in order to enhance our understanding of the human brain.

## Methods

### Participants

40 young and healthy adults were recruited from an internal participant database or through local advertisement and fully completed the experiment. No statistical methods were used to predetermine the sample size but it was chosen to be larger than similar previous neuroimaging studies [e.g., 49, 50, 52]. Four participants were excluded from further analysis because their mean behavioral performance was below the 50% chance level in either or both the sequence and repetition trials suggesting that they did not adequately process the visual stimuli used in the task. Thus, the final sample consisted of 36 participants (mean age = 24.61 years, *SD* = 3.77 years, age range: 20 - 35 years, 20 female, 16 male). All participants were screened for magnetic resonance imaging (MRI) eligibility during a telephone screening prior to participation and again at the beginning of each study session according to standard MRI safety guidelines (e.g., asking for metal implants, claustrophobia, etc.). None of the participants reported to have any major physical or mental health problems. All participants were required to be right-handed, to have corrected-to-normal vision, and to speak German fluently. Furthermore, only participants with a head circumference of 58 cm or less could be included in the study. This requirement was necessary as participants’ heads had to fit the MRI head coil together with MRI-compatible headphones that were used during the experimental tasks. The ethics commission of the German Psychological Society (DGPs) approved the study. All volunteers gave written informed consent prior to the beginning of the experiments. Every participant received 40.00 Euro and a performance-based bonus of up to 7.20 Euro upon completion of the study. None of the participants reported to have any prior experience with the stimuli or the behavioral task.

### Task

#### Stimuli

All stimuli were gray-scale images of a cat, chair, face, house, and shoe [cf. 53] with a size of 400 × 400 pixels each, which are freely available from http://data.pymvpa.org/datasets/haxby2001/ and have been shown to reliably elicit object-specific neural response patterns in several previous studies [e.g., 53, 56–58]. Participants received auditory feedback to signal the accuracy of their responses. A high-pitch coin sound confirmed correct responses, whereas a low-pitch buzzer sound signaled incorrect responses. The sounds were the same for all task conditions and were presented immediately after participants entered a response or after the response time had elapsed. Auditory feedback was used to anatomically separate the expected neural activation patterns of visual stimuli and auditory feedback. We recorded the presentation time stamps of all visual stimuli and confirmed that all experimental components were presented as expected. The task was programmed in MATLAB (version R2012b; Natick, Massachusetts, USA; The MathWorks Inc.) using the Psychophysics Toolbox extensions [version 3.0.11; 86–88] and run on a Windows XP computer with a monitor refresh-rate of 16.7 ms.

#### Slow trials

The slow trials of the task were designed to elicit object-specific neural response patterns of the presented visual stimuli. The resulting patterns of neural activation were later used to train the classifiers. In order to ensure that participants maintained their attention and processed the stimuli adequately, they were asked to perform an oddball detection task [for a similar approach, see 42, 45]. Specifically, participants were instructed to press a button each time an object was presented upside-down. Participants could answer using either the left or the right response button of an MRI-compatible button box. In contrast to similar approaches [e.g., 42, 45], we intentionally did not ask participants for a response on trials with upright stimuli to avoid neural activation patterns of motor regions in our training set which could influence later classification accuracy on the test set.

Participants were rewarded with 3 cents for each oddball (i.e., stimulus presented upside-down) that was correctly identified (i.e., hit) and punished with a deduction of 3 cents for (incorrect) responses (i.e., false alarms) on non-oddball trials (i.e., when stimuli were presented upright). In case participants missed an oddball (i.e., miss), they also missed out on the reward. Auditory feedback (coin and buzzer sound for correct and incorrect responses, respectively) was presented immediately after the response (in case of hits and false alarms) or at the end of the response time limit (in case of misses) using MRI-compatible headphones (VisuaStimDigital, Resonance Technology Company, Inc., Northridge, CA, USA). Correct rejections (i.e., no responses to upright stimuli) were not rewarded and were consequently not accompanied by auditory feedback. Together, participants could earn a maximum reward of 3.60 Euro in this task condition.

Across the entire experiment, all five unique images were presented in all possible sequential combinations which resulted in 5! = 120 sequences with each of the five unique visual objects in a different order. Thus, across the entire experiment participants were shown 120 ∗ 5 = 600 visual objects in total for this task condition. 20% of all visual objects were presented upside-down (i.e., 120 oddball stimuli). All unique visual objects were shown upside-down equally often, which resulted in 120/5 = 24 oddballs for each individual visual object category. The order of sequences as well as the appearances of oddballs were randomly shuffled for each participant and across both study sessions.

Each trial (for the trial procedure, see Fig. 1a) started with a waiting period of 3.85 s during which a blank screen was presented. This ITI ensured a sufficient time delay between each slow trial and the preceding trial (either a sequence or a repetition trial). The five visual object stimuli of the current trial were then presented as follows: After the presentation of a short fixation dot for a constant duration of 300 ms, a stimulus was shown for a fixed duration of 500 ms followed by a variable ISI during which a blank screen was presented again. The duration of the ISI for each trial was randomly drawn from a truncated exponential distribution with a mean of 2.5 s and a lower limit of 1 s. We expected that neural activation patterns elicited by the stimuli can be well recorded during this average time period of 3 s [for a similar approach, see 53]. Behavioral responses were collected during a fixed time period of 1.5 s after each stimulus onset. In case participants missed an oddball target, the buzzer sound (signaling an incorrect response) was presented after the response time limit had elapsed. Only neural activation patterns related to correct trials with upright stimuli were used to train the classifiers. Slow trials were interleaved with sequence and repetition trials such that each of the 120 slow trials was followed by either one of the 75 sequence trials or 45 repetition trials (details on these trial types follow below).

#### Sequence trials

On the sequence trials of the task, participants were shown sequences of the same five unique visual objects at varying presentation speeds. In total, 15 different sequences were selected for each participant. Sequences were chosen such that each visual object appeared equally often at the first and last position of the sequence. Given five stimuli and 15 sequences, for each object category this was the case for 3 out of the 15 sequences. Furthermore, we ensured that all possible sequences were chosen equally often across all participants. Given 120 possible sequential combinations in total, the sequences were distributed across eight groups of participants. Sequences were randomly assigned to each participant following this pseudo-randomized procedure.

To investigate the influence of sequence presentation speed on the corresponding neural activation patterns, we systematically varied the ISI between consecutive stimuli in the sequence. Specifically, we chose five different speed levels of 32, 64, 128, 512, and 2048 ms, respectively (i.e., all exponents of 2 for good coverage of faster speeds). Each of the 15 sequences per participant was shown at each of the 5 different speed levels. The occurrence of the sequences was randomly shuffled for each participant and across sessions within each participant. This resulted in a total of 75 sequence trials presented to each participant across the entire experiment. To ensure that participants maintained attention to the stimuli during the sequence trials, they were instructed to identify the serial position of a previously cued target object within the shown stimulus sequence and indicate their response after a delay period without visual input.

During a sequence trial (for the trial procedure, see Fig. 1b) the target cue (the name of the visual object, e.g., *shoe*) was shown for a fixed duration of 1000 ms, followed by a blank screen for a fixed duration of 3850 ms. A blank screen was used to reduce possible interference of neural activation patterns elicited by the target cue with neural response patterns following the sequence of visual objects. A short presentation of a gray fixation dot for a constant duration of 300 ms signaled the onset of the upcoming sequence of visual objects. All objects in the sequence were presented briefly for a fixed duration of 100 ms. The ISI for each trial was determined based on the current sequence speed (see details above) and was the same for all stimuli within a sequence. The sequence of stimuli was followed by a delay period with a gray fixation dot that was terminated once a fixed duration of 16 s since the onset of the first sequence object had elapsed. This was to ensure sufficient time to acquire the aftereffects of neural responses following the sequence of objects even at a sequence speed of 2048 ms. During the waiting period participants were listening to bird sounds (which can be downloaded from https://audiojungle.net/item/british-bird-song-dawn-chorus/98074) in order to keep them moderately entertained without additional visual input. Subsequently, the name of the target object as well as the response mapping was presented for a fixed duration of 1.5 s (same fixed response time limit as for the slow trials, see above). In this response interval, participants had to choose the correct serial position of the target object from two response options that were presented on the left and right side of the screen. The mapping of the response options was balanced for left and right responses (i.e., the correct option appeared equally often on the left and right side: 37 times each with the mapping of the last trial being determined randomly) and shuffled randomly for every participant. The serial position of the target for each trial was randomly drawn from a Poisson distribution with *λ* = 1.9 and truncated to an interval from 1 to 5. Thus, across all trials, the targets appeared more often at the later compared to earlier positions of the sequence. This was done to reduce the likelihood that participants stopped to process stimuli or diverted their attention after they identified the position of the target object. The serial position of the alternative response option was drawn from the same distribution as the serial position of the target. As for the oddball trials, auditory feedback was presented immediately following a response. The coin sound indicated a reward of 3 cents for correct responses, whereas the buzzer sound signaled incorrect or missed responses (however, there was no deduction of 3 cents for incorrect responses or misses). Together, participants could earn a maximum reward of 2.25 Euro in this task condition.

#### Repetition trials

We included so-called *repetition trials* to investigate how decoding time course would be affected by (1) the number of fast repetitions of the same neural event and (2) their interaction with the position of the switch to a subsequent stimulus category. Therefore, in this task condition, the same two stimuli were repeated a varying number of times each in one sequence. All sequences had a fixed length of nine stimuli in total. Each of the five stimulus categories was selected as the preceding stimulus for eight sequences in total. For each of these eight sequences we systematically varied the time point of the switch to the second stimulus category from serial position 2 to 9. Overall, the transition to the second stimulus happened five times at each serial position with varying stimulus material on each trial. Across the eight trials for each stimulus category, we ensured that each preceding stimulus category was followed by each of the remaining four stimulus categories equally often. Specifically, a given preceding stimulus category was followed by each of the remaining four stimulus categories two times. Also, the average serial position of the first occurrence of each of the subsequent stimuli was the same for all subsequent stimuli. That is to say, the same subsequent stimulus appeared either on position 9 and 2, 8 and 3, 7 and 4 or 6 and 5, resulting in an average first occurrence of the subsequent stimulus at position 5.5. All stimulus sequences of the repetition trials were presented with a fixed ISI of 32 ms. Note, that this is the same presentation speed as the fastest ISI of the sequence trials. Similar to the sequence trials, participants were instructed to remember the serial position at which the second stimulus within the sequence appeared for the first time. For example, if the switch to the second stimulus happened at the fifth serial position, participants had to remember this number.

Similar to the trial procedure of the sequence trials, each repetition trial (Fig. 1c) began with the presentation of the target cue (name of the visual object, e.g., *cat*), which was shown for a fixed duration of 500 ms. The target cue was followed by a blank screen that was presented for a fixed duration of 3.85 s. A briefly presented fixation dot announced the onset of the sequential visual stimuli. Subsequently, the fast sequence of visual stimuli was presented with a fixed duration for visual stimuli (100 ms each) and the ISI (32 ms on all trials). As for sequence trials, the sequence of stimuli on repetition trials was followed by a variable delay period until 16 s from sequence onset had elapsed. On repetition trials, participants had to choose the correct serial position of the first occurrence of the target stimulus from two response options. The incorrect response option was a random serial position that was at least two positions away from the correct target position. For example, if the correct option was 5, the alternative target position could either be earlier (1, 2, or 3) or later (7, 8, or 9). This was done to ensure that the task was reasonably easy to perform. Finally, we added five longer repetition trials with 16 elements per sequence. Here, the switch to the second sequential stimulus always occurred at the last serial position. Each of the five stimulus categories was the preceding stimulus once. The second stimulus of each sequence was any of the other four stimulus categories. In doing so, in the long repetition trials each stimulus category was the preceding and subsequent stimulus once. Repetition trials were randomly distributed across the entire experiment and (together with the sequence trials) interleaved with the slow trial.

### Study procedure

The study consisted of two experimental sessions. During the first session, participants were informed in detail about the study, screened for MRI eligibility, and provided written informed consent if they agreed to participate in the study. Then they completed a short demographic questionnaire (assessing age, education, etc.) and a computerized version of the Digit-Span Test, assessing working memory capacity [89]. Next, they performed a 10-minutes (min) practice of the main task. Subsequently, participants entered the MRI scanner. After a short localizer, we first acquired a 5-min resting state scan for which participants were asked to stay awake and focus on a white fixation cross presented centrally on a black screen. Then, we acquired four functional task runs of about 11 min during which participants performed the main task in the MRI scanner. After the functional runs, we acquired another 5-min resting state, 5-min fieldmaps as well as a 4-min anatomical scan. The second study session was identical to the first session, except that participants entered the scanner immediately after another short assessment of MRI eligibility. In total, the study took about four hours to complete (2.5 and 1.5 hours for Session 1 and 2, respectively).

### MRI data acquisition

All MRI data were acquired using a 32-channel head coil on a research-dedicated 3-Tesla Siemens Magnetom TrioTim MRI scanner (Siemens, Erlangen, Germany) located at the Max Planck Institute for Human Development in Berlin, Germany. The scanning procedure was exactly the same for both study sessions. For the functional scans, whole-brain images were acquired using a segmented k-space and steady state T2*-weighted multi-band (MB) echo-planar imaging (EPI) single-echo gradient sequence that is sensitive to the BOLD contrast. This measures local magnetic changes caused by changes in blood oxygenation that accompany neural activity (sequence specification: 64 slices in interleaved ascending order; anterior-to-posterior (A-P) phase encoding direction; TR = 1250 ms; echo time (TE) = 26 ms; voxel size = 2 × 2 × 2 mm; matrix = 96 × 96; field of view (FOV) = 192 × 192 mm; flip angle (FA) = 71 degrees; distance factor = 0%; MB acceleration factor 4). Slices were tilted for each participant by 15 degrees forwards relative to the rostro-caudal axis to improve the quality of fMRI signal from the hippocampus [cf. 90] while preserving good coverage of occipito-temporal brain regions. Each MRI session included four functional task runs. Each run was about 11 minutes in length, during which 530 functional volumes were acquired. For each functional run, the task began after the acquisition of the first four volumes (i.e., after 5.00 s) to avoid partial saturation effects and allow for scanner equilibrium. We also recorded two functional runs of resting-state fMRI data, one before and one after the task runs. Each resting-state run was about 5 minutes in length, during which 233 functional volumes were acquired. After the functional task runs, two short acquisitions with six volumes each were collected using the same sequence parameters as for the functional scans but with varying phase encoding polarities, resulting in pairs of images with distortions going in opposite directions between the two acquisitions (also known as the *blip-up / blip-down* technique). From these pairs the displacements map were estimated and used to correct for geometric distortions due to susceptibility-induced field inhomogeneities as implemented in the the fMRIPrep preprocessing pipeline [91]. In addition, a whole-brain spoiled gradient recalled (GR) field map with dual echo-time images (sequence specification: 36 slices; A-P phase encoding direction; TR = 400 ms; TE1 = 4.92 ms; TE2 = 7.38 ms; FA = 60 degrees; matrix size = 64 × 64; FOV = 192 × 192 mm; voxel size = 3 × 3 × 3.75 mm) was obtained as a potential alternative to the method described above. However, as this field map data was not successfully recorded for four participants, we used the blip-up blip-down technique for distortion correction (see details on MRI data pre-processing below). Finally, high-resolution T1-weighted (T1w) anatomical Magnetization Prepared Rapid Gradient Echo (MPRAGE) sequences were obtained from each participant to allow registration and brain surface reconstruction (sequence specification: 256 slices; TR = 1900 ms; TE = 2.52 ms; FA = 9 degrees; inversion time (TI) = 900 ms; matrix size = 192 × 256; FOV = 192 × 256 mm; voxel size = 1 × 1 × 1 mm). We also measured respiration and pulse during each scanning session using pulse oximetry and a pneumatic respiration belt.

### MRI data preparation and preprocessing

Results included in this manuscript come from preprocessing performed using *fMRIPrep* 1.2.1 (Esteban et al. [91, 92]; RRID:SCR 016216), which is based on *Nipype* 1.1.4 (Gorgolewski et al. [93, 94]; RRID:SCR 002502). Many internal operations of *fMRIPrep* use *Nilearn* 0.4.2 [95, RRID:SCR 001362], mostly within the functional processing workflow. For more details of the pipeline, see the section corresponding to workflows in *fMRIPrep*’s documentation.

#### Conversion of data to the brain imaging data structure (BIDS) standard

The majority of the steps involved in preparing and preprocessing the MRI data employed recently developed tools and workflows aimed at enhancing standardization and reproducibility of task-based fMRI studies [for a similar preprocessing pipeline, see 96]. Following successful acquisition, all study data were arranged according to the BIDS specification [97] using the HeuDiConv tool (version 0.6.0.dev1; freely available from https://github.com/nipy/heudiconv) running inside a Singularity container [98, 99] to facilitate further analysis and sharing of the data. Dicoms were converted to the NIfTI-1 format using dcm2niix [version 1.0.20190410 GCC6.3.0; 100]. In order to make identification of study participants unlikely, we eliminated facial features from all high-resolution structural images using pydeface (version 2.0; available from https://github.com/poldracklab/pydeface). The data quality of all functional and structural acquisitions were evaluated using the automated quality assessment tool MRIQC [for details, see 101, and the MRIQC documentation]. The visual group-level reports of the estimated image quality metrics confirmed that the overall MRI signal quality of both anatomical and functional scans was highly consistent across participants and runs within each participant.

#### Preprocessing of anatomical MRI data

A total of two T1w images were found within the input BIDS data set, one from each study session. All of them were corrected for intensity non-uniformity (INU) using N4BiasFieldCorrection [Advanced Normalization Tools (ANTs) 2.2.0; 102]. A T1w-reference map was computed after registration of two T1w images (after INU-correction) using mri robust template [FreeSurfer 6.0.1, 103]. The T1w-reference was then skull-stripped using antsBrainExtraction.sh (ANTs 2.2.0), using OASIS as target template. Brain surfaces were reconstructed using recon-all [FreeSurfer 6.0.1, RRID:SCR 001847, 104], and the brain mask estimated previously was refined with a custom variation of the method to reconcile ANTs-derived and FreeSurfer-derived segmentations of the cortical gray-matter of Mindboggle [RRID:SCR 002438, 105]. Spatial normalization to the ICBM 152 Nonlinear Asymmetrical template version 2009c [106, RRID:SCR 008796] was performed through nonlinear registration with antsRegistration [ANTs 2.2.0, RRID:SCR 004757, 107], using brain-extracted versions of both T1w volume and template. Brain tissue segmentation of cerebrospinal fluid (CSF), white-matter (WM) and gray-matter (GM) was performed on the brain-extracted T1w using fast [FSL 5.0.9, RRID:SCR 002823, 108].

#### Preprocessing of functional MRI data

For each of the BOLD runs found per participant (across all tasks and sessions), the following preprocessing was performed. First, a reference volume and its skull-stripped version were generated using a custom methodology of *fMRIPrep*. The BOLD reference was then co-registered to the T1w reference using bbregister (FreeSurfer) which implements boundary-based registration [109]. Co-registration was configured with nine degrees of freedom to account for distortions remaining in the BOLD reference. Head-motion parameters with respect to the BOLD reference (transformation matrices, and six corresponding rotation and translation parameters) are estimated before any spatiotemporal filtering using mcflirt [FSL 5.0.9, 110]. BOLD runs were slice-time corrected using 3dTshift from AFNI 20160207 [111, RRID:SCR 005927]. The BOLD time-series (including slice-timing correction when applied) were resampled onto their original, native space by applying a single, composite transform to correct for head-motion and susceptibility distortions. These resampled BOLD time-series will be referred to as *preprocessed BOLD in original space*, or just *preprocessed BOLD*. The BOLD time-series were resampled to MNI152NLin2009cAsym standard space, generating a *preprocessed BOLD run in MNI152NLin2009cAsym space*. First, a reference volume and its skull-stripped version were generated using a custom methodology of *fMRIPrep*. Several confounding time-series were calculated based on the *preprocessed BOLD*: framewise displacement (FD), DVARS and three region-wise global signals. FD and DVARS are calculated for each functional run, both using their implementations in *Nipype* [following the definitions by 112]. The three global signals are extracted within the CSF, the WM, and the whole-brain masks. Additionally, a set of physiological regressors were extracted to allow for component-based noise correction [*CompCor*, 113]. Principal components are estimated after high-pass filtering the *preprocessed BOLD* time-series (using a discrete cosine filter with 128s cut-off) for the two *CompCor* variants: temporal (tCompCor) and anatomical (aCompCor). Six tCompCor components are then calculated from the top 5% variable voxels within a mask covering the subcortical regions. This subcortical mask is obtained by heavily eroding the brain mask, which ensures it does not include cortical GM regions. For aCompCor, six components are calculated within the intersection of the aforementioned mask and the union of CSF and WM masks calculated in T1w space, after their projection to the native space of each functional run (using the inverse BOLD-to-T1w transformation). The head-motion estimates calculated in the correction step were also placed within the corresponding confounds file. The BOLD time-series, were resampled to surfaces on the following spaces: *fsnative, fsaverage*. All resamplings can be performed with *a single interpolation step* by composing all the pertinent transformations (i.e., head-motion transform matrices, susceptibility distortion correction when available, and co-registrations to anatomical and template spaces). Gridded (volumetric) resamplings were performed using antsApplyTransforms (ANTs), configured with Lanczos interpolation to minimize the smoothing effects of other kernels [114]. Non-gridded (surface) resamplings were performed using mri vol2surf (FreeSurfer). Following preprocessing using fMRIPrep, the fMRI data were spatially smoothed using a Gaussian mask with a standard deviation (Full Width at Half Maximum (FWHM) parameter) set to 4 mm using an example Nipype smoothing workflow (see the Nipype documentation for details) based on the SUSAN algorithm as implemented in the FMRIB Software Library (FSL) [115].

### Multi-variate fMRI pattern analysis

#### Leave-one-run-out cross-validation procedure

All fMRI pattern classification analyses were conducted using open-source packages from the Python (Python Software Foundation, Python Language Reference, version 3.7) modules Nilearn [version 0.5.0; 95] and scikit-learn [version 0.20.3; 116]. fMRI pattern classification was performed using a leave-one-run-out cross-validation procedure for which data from seven task runs were used for training and data from the left-out run (i.e., the eighth run) was used for testing. This procedure was repeated eight times so that each task run served as the training set once. We trained an ensemble of five independent classifiers, one for each of the five stimulus classes (cat, chair, face, house, and shoe). For each class-specific classifier, labels of all other classes in the data were relabelled to a common *other* category. In order to ensure that the classifier estimates were not biased by relative differences in class frequency in the training set, the weights associated with each class were adjusted inversely proportional to the class frequencies in each training fold. Training was performed on data from all trials of the seven runs in the respective cross-validation fold only using trials of the slow task where the visual object stimuli were presented upright and participants correctly did not respond (i.e., correct rejection trials). In each iteration of the classification procedure, the classifiers trained on seven out of eight runs were then applied separately to the data from the left-out run. Specifically, the classifiers were applied to (1) data from the slow trials of the left-out run, selecting volumes capturing the expected activation peaks to determine classification accuracy, (2) data from the slow trials of the left-out run, selecting all volumes from stimulus onset to the end of the trial (seven volumes in total per trial) to identify temporal dynamics of classifier predictions on a single trial basis, (3) data from the sequence trials of the left-out run, selecting all volumes from sequence onset to the end of the delay period (13 volumes in total per trial), (4) data from the repetition trials of the left-out run, also selecting all volumes from sequence onset to the end of the delay period (13 volumes in total per trial).

We used separate multinomial logistic regression classifiers with identical parameter settings. All classifiers were regularized using L2 regularization. The *C* parameter of the cost function was fixed at the default value of 1.0 for all participants. The classifiers employed the lbfgs algorithm to solve the multi-class optimization problem and were allowed to take a maximum of 4, 000 iterations to converge. Pattern classification was performed within each participant separately, never across participants. For each stimulus in the training set, we added 4 s to the stimulus onset and chose the volume closest to that time point (i.e., rounded to the nearest volume) to center the classifier training on the expected peaks of the BOLD response [for a similar approach, see e.g., 47]. At a TR of 1.25 s this corresponded to the fourth MRI volume which thus compromised a time window of 3.75 s to 5 s after each stimulus onset. We detrended the fMRI data separately for each run across all task conditions to remove low frequency signal intensity drifts in the data due to noise from the MRI scanner. For each classifier and run, the features were standardized (z-scored) by removing the mean and scaling to unit variance separately for each test set.

For fMRI pattern classification analysis performed on resting-state data we created a new mask for each participant through additive combination of the eight masks used for cross-validation (see above). This mask was then applied to all task and resting-state fMRI runs which were then separately detrended and standardized (z-scored). The classifiers were trained on the peak activation patterns from all slow trials combined.

#### Feature selection

Feature selection is commonly used in multi-voxel pattern analysis (MVPA) to determine the voxels constituting the activation patterns used for classification in order to improve the predictive performance of the classifier [117, 118]. Here, we combined a functional ROI approach based on thresholded *t*-maps with anatomical masks to select image-responsive voxels within a predefined anatomical brain region.

We ran eight standard first-level general linear models (GLMs) for each participant, one for each of the eight cross-validation folds using SPM12 (version 12.7219; https://www.fil.ion.ucl.ac.uk/spm/software/spm12/) running inside a Singularity container built using neurodocker (https://github.com/ReproNim/neurodocker) implemented in a custom analysis workflow using Nipype [version 1.4.0; 93]. In each cross-validation fold, we fitted a first-level GLM to the data in the training set (e.g., data from run 1 to 7) and modeled the stimulus onset of all trials of the slow task when a stimulus was presented upright and was correctly rejected (i.e., participants correctly did not respond). These trial events were modeled as boxcar functions with the length of the modeling event corresponding to the duration of the stimulus on the screen (500 ms for all events). If present in the training data, we also included trials with hits (correct response to upside-down stimuli), misses (missed response to upside-down stimuli) and false alarms (incorrect response to upright stimuli) as regressors of no interest, thereby explicitly modeling variance attributed to these trial types [cf. 119]. Finally, we included the following nuisance regressors estimated during preprocessing with fMRIPrep: the frame-wise displacement for each volume as a quantification of the estimated bulk-head motion, the six rigid-body motion-correction parameters estimated during realignment (three translation and rotation parameters, respectively), and six noise components calculated according to the anatomical variant of *CompCorr* [for details, see 91, and the fMRIPrep documentation]. All regressors were convolved with a canonical HRF and did not include model derivatives for time and dispersion. Serial correlations in the fMRI time series were accounted for using an autoregressive AR(1) model. This procedure resulted in fold-specific maps of *t*-values that were used to select voxels from the left-out run of the cross-validation procedure. Note, that this approach avoids circularity (or so-called *double-dipping*) as the selective analysis (here, fitting of the GLMs to the training set) is based on data that is fully independent from the data that voxels are later selected from [here, testing set from the left-out run; cf. 120].

The resulting brain maps of voxel-specific *t*-values resulting from the estimation of the described *t*-contrast were then combined with an anatomical mask of occipito-temporal brain regions. All participant-specific anatomical masks were created based on automated anatomical labeling of brain surface reconstructions from the individual T1w reference image created with Freesurfer’s recon-all [104] as part of the fMRIPrep workflow [91], in order to account for individual variability in macroscopic anatomy and to allow reliable labeling [121, 122]. For the anatomical masks of occipito-temporal regions we selected the corresponding labels of the cuneus, lateral occipital sulcus, pericalcarine gyrus, superior parietal lobule, lingual gyrus, inferior parietal lobule, fusiform gyrus, inferior temporal gyrus, parahippocampal gyrus, and the middle temporal gyrus [cf. 53]. Only gray-matter voxels were included in the generation of the masks as BOLD signal from non-gray-matter voxels cannot be generally interpreted as neural activity [118]. Note, however, that due to the whole-brain smoothing performed during preprocessing, voxel activation from brain regions outside the anatomical mask but within the sphere of the smoothing kernel might have entered the anatomical mask (thus, in principle, also including signal from surrounding non-gray-matter voxels).

Finally, we combined the *t*-maps derived in each cross-validation fold with the anatomical masks. All voxels with *t*-values above or below a threshold of *t* = 3 (i.e., voxels with the most negative and most positive *t*-values) inside the anatomical mask were then selected for the left-out run of the classification analysis and set to 1 to create the final binarized masks (*M* = 11162 voxels on average, *SD* = 2083).

#### Classification accuracy and multivariate decoding time courses

In order to assess the classifiers’ ability to differentiate between the neural activation patterns of individual visual objects, we compared the predicted visual object of each example in the test set to the visual object that was actually shown to the participant on the corresponding trial. We obtained an average classification accuracy score for each participant by calculating the mean proportion of correct classifier predictions across all correctly answered, upright slow trials (Fig. 2a). The mean accuracy scores of all participants were then compared to the chance baseline of 100%/5 = 20% using a one-sided one-sample t-test, testing the a-priori hypothesis that classification accuracy would be higher than the chance baseline. The effect size (Cohen’s *d*) was calculated as the difference between the mean of accuracy scores and the chance baseline, divided by the standard deviation of the data [123]. Furthermore, we assessed the classifiers’ ability to accurately detect the presence of visual objects on a single trial basis. For this analysis we applied the trained classifiers to seven volumes from the volume closest to the stimulus onset, which allowed us to examine the time courses of the probabilistic classification evidence in response to the visual stimuli on a single trial basis (Fig. 2b). In order to test if the time series of classifier probabilities reflected the expected increase of classifier probability for the stimulus shown on a given trial, we compared the time series of classifier probabilities related to the classified class with the mean time courses of all other classes using a two-sided paired t-test at every time point (i.e., at every TR). Here, we used the Bonferroni-correction method [124] across time points and stimulus classes to adjust for multiple comparisons of 35 observations (7 TRs and 5 stimulus classes). In the main text, we only report the results for the peak in classification probability of the true class, corresponding to the fourth TR after stimulus onset. The effect size (Cohen’s *d*) was calculated as the difference between the means of the probabilities of the current versus all other stimuli, divided by the standard deviation of the difference [123].

#### Response and difference function modeling

As reported above, analyzing probabilistic classifier evidence on single slow trials revealed multivariate decoding time courses that can be characterized by a slow response function that resembles single-voxel hemodynamics. For simplicity, we modelled this response function as a sine wave that was flattened after one cycle, scaled by an amplitude and adjusted to baseline. The model was specified as follows:

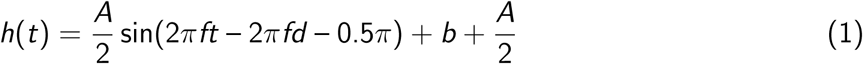

whereby *A* is the response amplitude (the peak deviation of the function from baseline), *f* is the angular frequency (unit: 1/TR, i.e., 0.8 Hz), *d* is the onset delay (in TRs), and *b* is the baseline (in %). The restriction to one cycle was achieved by converting the sine wave in accordance with the following piecewise function

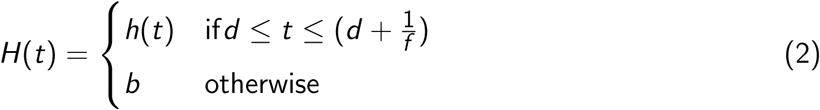

We fitted the four model parameters (*A, f, d* and *b*) to the mean probabilistic classifier evidence of each stimulus class at every TR separately for each participant. For convenience, we count time *t* in TRs. To approximate the time course of the difference between two response functions we utilized the trigonometric identity for the subtraction of two sine functions [e.g., 125]:

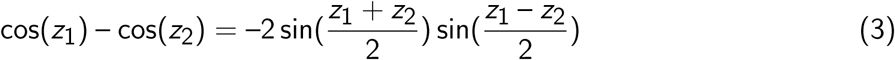

Considering the case of two sine waves with identical frequency but differing by a temporal shift *δ* one obtains

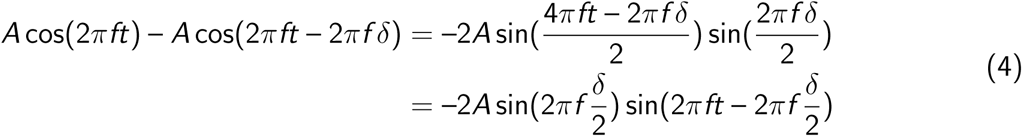

which corresponds to a flipped sine function with an amplitude scaled by 2 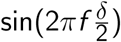, a shift of 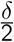 and an identical frequency *f* .

To apply this equation to our scenario two adjustments have to be made since the the single-cycle nature of our response function is not accounted for in Equation 3. First, one should note that properties of the amplitude term in Equation 4 only hold as long as shifts of no greater than half a wavelength are considered (the wavelength *λ* is the inverse of the frequency *f*). The term 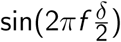 can be written as 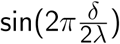, which illustrates that the term monotonically increases until 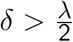. Second, the frequency term has to be adapted as follows: The flattening of the sine waves to the left implies that the difference becomes positive at 0 rather than 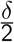, thus undoing the phase shift and stretching the wave by 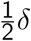 TRs. The flattening on the right also leads to a lengthening of the wave by an additional 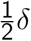 TRs, since the difference becomes 0 at 2*πf* + 2*πf δ*, instead of only 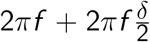. Thus, the total wavelength has to be adjusted by a factor of *δ* TRs, and no phase shift relative to the first response is expected. The difference function therefore has frequency

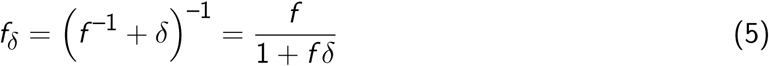

instead of *f*, and Equation 4 becomes 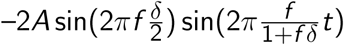. We can now apply Equation 3 to the fitted response function as follows

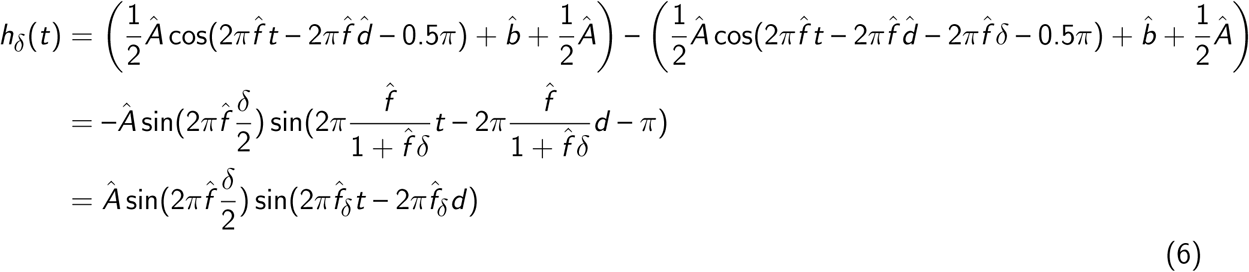

whereby 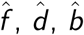 and *Â* indicate fitted parameters.

We determined the relevant TRs in the forward and backward periods for sequence trials by calculating *δ* depending on the sequence speed (the ISI). The resulting values for *δ* and corresponding forward and backward periods are shown in Table 1. Model fitting was performed using NLoptr, an R interface to the NLopt library for nonlinear optimization [126] employing the COBYLA (Constrained Optimization BY Linear Approximation) algorithm [127, 128]. The resulting parameters were then averaged across participants, yielding the mean parameters reported in the main text. To assess if the model fitted the data reasonably, we inspected the fits of the sine wave response function for each stimulus class and participant using individual parameters (Fig. S2).

**Table 1:** Relevant time periods depending on sequence speed. Forward periods were calculated as [0.56; 0.5 ∗ *λ*_*δ*_ + *d* = 0.5 ∗ (5.26 + *δ*) + 0.56]. Backward period were calculated as [0.5 ∗ *λ*_*δ*_ + *d* = 0.5 ∗ (5.26 + *δ*) + 0.56; *λ*_*δ*_ + *d* = 5.26 + *δ* + 0.56]. *δ* reflects the interval between the onsets of the first and last of five sequence items that is dependent on the sequence speed (the ISI) and the stimulus duration (here, 100 ms). For example, for an ISI of 32 ms, *δ* (in TRs) is calculated as (0.032 ∗ 4 + 0.1 ∗ 4)/1.25 = 0.42 TRs. *d* reflects the fitted onset delay (here, 0.56 TRs). All values were then rounded to the closest TRs resulting in the speed-adjusted time periods (two rightmost columns).

| Speed | $\delta$ (in TRs) | Forward period | Backward period |
| --- | --- | --- | --- |
| 32 ms | 0.42 TRs | TRs 2–4 | TRs 5–7 |
| 64 ms | 0.52 TRs | TRs 2–4 | TRs 5–7 |
| 128 ms | 0.73 TRs | TRs 2–4 | TRs 5–8 |
| 512 ms | 1.96 TRs | TRs 2–5 | TRs 6–9 |
| 2048 ms | 6.87 TRs | TRs 2–7 | TRs 8–13 |

#### Detecting sequentiality in fMRI patterns on sequence trials

In order to analyze the neural activation patterns following the presentation of sequential visual stimuli for evidence of sequentiality, we first determined the true serial position of each decoded event for each trial. Specifically, applying the trained classifiers to each volume of the sequence trials yielded a series of predicted event labels and corresponding classification probabilities that were assigned their sequential position within the true sequence that was shown to participants on the corresponding trial.

The main question we asked for this analysis was to what extend we can infer the serial order of image sequences from relative activation differences in fMRI pattern strength within single measurements (a single TR). To this end, we applied the trained classifiers to a series of 13 volumes following sequence onset (spanning a total time window of about 16 s) on sequence trials and analyzed the time courses of the corresponding classifier probabilities related to the five image categories (Fig. 3a). Classification probabilities were normalized by dividing the probabilities by their trial-wise sum for each image class. As detailed in the task description, the time window was selected such that the neural responses to the image sequences could be fully captured without interference from upcoming trials. We examined relative differences in decoding probabilities between serial events at every time-point (i.e., at every TR) and quantified the degree of sequential ordering in two different analyses:

First, we conducted a linear regression between the serial position of the five images and their classification probabilities at every TR in the relevant forward and backward period (adjusted by sequence speed) and extracted the slope of the linear regression as an index of linear association. The slopes were then averaged at every TR separately for each participant and sequence speed across data from all fifteen sequence trials (Fig. 3b). Here, if later events have a higher classification probability compared to earlier events, the slope coefficient will be negative. In contrast, if earlier events have a higher classification probability compared to later events, the slope coefficient will be positive. Note, that for convenience, we flipped the sign of the mean regression slopes so that positive values indicate forward ordering and negative values indicate backward ordering. To determine if we can find evidence for significant sequential ordering of classification probabilities in the forward and backward periods, we conducted a series of ten separate two-tailed one-sample t-tests comparing the mean regression slope coefficients of each speed condition against zero (the expectation of no order information). All *p* values were adjusted for ten comparisons by controlling the FDR (Fig. 3c; [129]). As an estimate of the effect size, we calculated Cohen’s *d* as the difference between the sample mean and the null value in units of the sample standard deviation [123]. As reported in the main text, we conducted the same analysis using ranked correlation coefficients (Kendall’s *τ*) and the mean step size between probability-ordered events within TRs as alternative indices of linear association (for details, see SI). In order to directly compare the predicted time courses of regression slopes based on our modeling approach with the observed time courses, we computed the Pearson’s correlation coefficient between the two time series both on data averaged across participants and within each participant (Figs. 2d–e). The mean within-participant correlation coefficients were tested against zero (the expectation of no correlation) using a separate two-sided one-sample t-test for each speed condition. All *p* values were adjusted for five comparisons by controlling the FDR [129]).

We hypothesized that sequential order information of fast neural events will translate into order structure in the fMRI signal and successively decoded events in turn. Therefore, we analyzed the fMRI data from sequence trials for evidence of sequentiality across consecutive measurements. The analyses were restricted to the expected forward and backward periods which were adjusted depending on the sequence speed. For each TR we obtained the image with the most likely fMRI signal pattern based on the classification probabilities. First, we asked if we are more likely to decode earlier serial events earlier and later serial events later in the decoding time window of thirteen TRs. To this end, we averaged the serial position of the most likely event at every TR, separately for each trial and participant, resulting in a time course of average serial event position across the decoding time window (Fig. 3d). We then compared the average serial event position against the mean serial position (position 3) as a baseline across participants at every time point in the forward and backward period using a series of two-sided one-sample t-tests, adjusted for 38 multiple comparisons (across all five speed conditions and TRs in the forward and backward period) by controlling the FDR [129]. These results are reported in the SI. Next, in order to assess if the average serial position differed between the forward and backward period for the five different speed conditions, we conducted a linear mixed effects (LME) and entered the speed condition (with five levels) and trial period (forward versus backward) as fixed effects including by-participant random intercepts and slopes. Finally, we conducted a series of two-sided one-sample t-tests to assess whether the mean serial position in the forward and backward periods differed from the expected mean serial position (baseline of 3) for every speed condition (all *p* values adjusted for 10 comparisons using FDR correction [129]).

Second, we analyzed how this progression through the involved sequence elements affected transitions between consecutively decoded serial events. As before, we extracted the most likely pattern for each TR (i.e., the pattern with the highest classification probability), and calculated the step sizes between consecutively decoded serial events, as in [52]. For example, decoding Event 2 → Event 4 in consecutive TRs would correspond to a step size of +2, while a Event 3 → Event 2 transition would reflect a step size of –1, etc. We then calculated the mean step-size of the first (early) and second (late) halves of the forward and backward periods, respectively, which were adjusted for sequence speed. Specifically, the transitions were defined as follows: at speeds of 32, 64 and 128 ms these transitions included the 2 → 3 (early forward), 3 → 4 (late forward), 5 → 6 (early backward) and 6 → 7 (late backward); at speeds of 512 ms these transitions included 2 → 3 (early forward), 4 → 5 (late forward), 6 → 7 (early backward), and 8 → 9 (late backward); at 2048 ms these transitions included 2 → 3 → 4 (early forward), 5 → 6 → 7 (late backward) 8 → 9 → 10 (early backward), and 11 → 12 → 13 (late backward). Finally, we compared the mean step size in the early and late half of the forward versus backward period for every speed condition using ten separate two-sided one-sample t-tests. All *p*s were adjusted for multiple comparisons by controlling the FDR [cf. 129].

#### Analysis of repetition trials for sensitivity of within-sequence items

Applying the classifiers trained on slow trials to data from *repetition trials* yielded a classification probability estimate for each stimulus class given the data at every time point (i.e., at every TR, Fig. 4a, S7). As described in the main text, we then analyzed the classification probabilities to answer which fMRI pattern were activated during a fast sequence under conditions of extreme forward or backward interference. Specifically, sequences with forward interference entailed a brief presentation of a single image that was followed by eight repetitions of a second image; whereas backward interference was characterized by a condition where eight image repetitions were followed by a single briefly presented item. As predicted by the sine-based response functions, the relevant time period included TRs 2–7. All analyses reported in the Results section were conducted using data from these selected TRs as described. Results based on data from all TRs are reported in the SI.

First, we calculated the mean probability of each event type (*first, second*, and *non-sequence* events) across all selected TRs and trials in the relevant time period separately for each repetition condition across participants. In order to examine whether the event type (*first, second*, and *non-sequence* events) had an influence on the mean probability estimates on *repetition trials*, we conducted a LME model [130] and entered the event type (with three factor levels: *first, second*, and *non-sequence* events) as a fixed effect and included by-participant random intercepts and slopes (Fig. 4b). Post-hoc comparisons between the means of the three factor levels were conducted using Tukey’s honest significant difference (HSD) test [131].

Second, in order to jointly examine the influence of event duration (number of repetitions) and event type (*first, second*, and *non-sequence* events), we conducted a LME model [130] with fixed effects of event type (with three factor levels: *first, second*, and *non-sequence* events) and repetition condition (number of individual event repetitions with two factor levels: (1) *forward interference* trials, where one briefly presented event is followed by eight repetitions of a second event, and (2) *backward interference* trials, where eight repetitions of a first event are followed by one briefly presented second event), also adding an interaction term for the two effects. Again, the model included both by-participant random intercepts and slopes (Fig. 4c). Post-hoc multiple comparisons among interacting factor levels were performed separately for each repetition condition by conditioning on each level of this factor (i.e., forward interference versus backward interference trials), using Tukey’s HSD test.

Third, we asked if we are more likely to find transitions between decoded events that were part of the sequence (the two within-sequence items) compared to items that were not part of the sequence (non-sequence items). To this end, we classified each transition as follows: forward (from Event 1 to Event 2), backward (from Event 2 to Event 1), repetitions of each sequence item, outwards (from sequence items to any non-sequence item), inwards (from non-sequence items to sequence items), outside (among non-sequence items) and repetitions among non-sequence events (the full transition matrix is shown in Fig. 4e). We then compared the average proportion of forward transitions within the sequence (i.e., decoding a Event 1 → Event 2) with the average proportions of (1) transitions from sequence items to items that were not part of the sequence (outwards transitions), and (2) transitions between events not part of the sequence (outside transitions) using paired two-sample t-tests with *p*s adjusted for four comparisons using Bonferroni correction (Fig. 4d).

#### Analysis of sparse sequence events with lower SNR

We only used resting state data from the first study session before participants had any experience with the task (except a short training session outside the scanner). These resting state data could not be successfully recorded in four participants. Therefore, the analyses were restricted to *N* = 32 of 36 participants. Participants were instructed to rest as calmly as possible with eyes opened while focusing on a white fixation cross that was presented centrally on the screen. For decoding on resting state data, we used the union of all eight masks created for the functional task runs during the cross-validation procedure. Logistic regression classifiers were trained on masked data from slow trials of all eight functional runs and applied to all TRs of the resting state data, similar to our sequence trial analysis. We assigned pseudo serial positions to each class randomly for every participant, assuming one fixed event ordering. We first characterized and compared the behavior of sequence detection metrics on resting state and concatenated sequence trial data. For sequence trials, we only considered data from TRs within the expected forward and backward periods (TRs 2 to 13) and focused on the fastest (32 ms) and slowest (2048 ms) speed condition. Accordingly, we restricted the resting state data to the first 180 TRs to match it to the length of concatenated sequence trial data (15 concatenated trials of 12 TRs each). For both fast and slow sequence trials and rest data, we then calculated the standard deviation of the probabilities (Fig. 5a) as well as the slope of a linear regression between serial position and their classification probabilities (Fig. 5b, 5c) at every TR. We then compared both the standard deviation of probabilities and the mean regression slopes over the entire rest period with the mean regression slopes in fast (32 ms) sequence trials using two-sided paired t-tests (Fig. 5a, 5b). *p*s adjusted for four comparisons using Bonferroni correction (Fig. 4d). The effect sizes (Cohen’s *d*) were calculated as the difference between the means of the resting and sequence data, divided by the standard deviation of the differences [123]. Given the rhythmic fluctuations of the regression slope dynamics (Fig. 2e) we calculated the frequency spectra across the resting state and concatenated sequence trial data using the Lomb-Scargle method [using the lsp function from the R package lomb, e.g., 60] that is suitable for unevenly-sampled data and therefore accounts for potential artifacts due to data concatenation 5d). The resulting frequency spectra were smoothed with a running average filter with width 0.005. Next, we extracted the mean power of the frequencies for fast and slow event sequences as predicted by Eqn. 5 in both resting and sequence data. For example, for a 32 ms sequence with *δ* = 0.032 ∗ 4 + 0.1 ∗ 5 = 0.628 one obtains the predicted frequency as 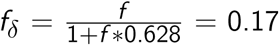, whereby *f* equals the fitted single trial frequency *f* = 1/5.26. The mean power at the predicted frequencies were then compared between resting as well as fast and slow sequence data using two-sided paired t-tests with *p* values adjusted for multiple comparisons using FDR-correction [129].

We then inserted 1 to 6 sequence events into the pre-task resting state period by blending TRs during resting state with TRs recorded during fast (32 ms) or slow (2048 ms) sequence trials. Specifically, we randomly selected six sequence trials for each speed condition, without replacement. Only TRs from the relevant time period (see above; 12 TRs for both speed conditions, respectively) were blended into the resting state data. To investigate the effects of a reduced SNR we systematically multiplied the probabilities of the inserted sequence TRs by a factor *κ* of 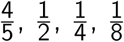 or 0, step-wise reducing the signal from 80% to 0% and added these scaled probabilities to the probability time courses of the resting state data. The resting state data used for blending were independently sampled from non-overlapping random locations within the resting state data of the same participant. This ensured that even in the 0 SNR condition, potential artefacts due to data concatenation were present and would therefore not impact our comparisons between SNR levels. For each combination of the number of inserts and SNR levels, we then compared the mean standard deviation of the probabilities during sequence-inserted rest with sequence-free rest using a series of two-sided paired t-tests. *p* values were adjusted accordingly for 30 comparisons using FDR-correction [129] and log-transformed (base 20) to make them easier to visualize (here, a log-transformed *p* values of 1 corresponds to *p* < .05).

Finally, we calculated the frequency spectra of sequence-inserted rest data as before, separately for data with fast and slow sequence inserts. To achieve comparable resolution obtained in the above analyses, we over-sampled the frequency space by a factor of 2. Smoothing was then applied again as before. We then calculated the relative power of each frequency compared to sequence-free rest and averaged the relative frequency spectra across participants (Fig. 5h). As before, we extracted the mean power within the predicted fast and slow frequency range (±0.01 Hz, given the smoothing) and compared them between fast and slow sequence-inserted rest and for different numbers of inserts and SNR levels. We them compared the relative power for each sequence-inserted rest data set, number of inserts and SNR level against zero (no difference from sequence-free rest) using a series of two-sided one-sample t-tests (*p* values uncorrected).

#### Statistical analysis

Main statistical analyses were conducted using LME models employing the lmer function of the lme4 package [version 1.1.21, 130] in R [version 3.6.1, 132]. If not stated otherwise, all models were fit with participants considered as a random effect on both the intercept and slopes of the fixed effects, in accordance with results from Barr et al. [133] who recommend to fit the most complex model consistent with the experimental design [133]. If applicable, explanatory variables were standardized to a mean of zero and a standard deviation of one before they entered the models. If necessary, we removed by-participant slopes from the random effects structure to allow a non-singular fit of the model [133]. Models were fitted using the BOBYQA (Bound Optimization BY Quadratic Approximation) optimizer [134, 135] with a maximum of 500, 000 function evaluations and no calculation of gradient and Hessian of nonlinear optimization solution. The likelihoods of the fitted models were assessed using Type III analysis of variance (ANOVA) with Satterthwaite’s method. A single-step multiple comparison procedure between the means of the relevant factor levels was conducted using Tukey’s HSD test [131], as implemented in the emmeans package in R [version 1.3.4, 132, 136]. In all other analyses we used one-sample t-tests if group data was compared to e.g., a baseline or paired t-tests if two sample from the same population were compared. If applicable, correction for multiple hypothesis testing was performed using the FDR-correction method [129]. If not stated otherwise, t-tests were two-sided and the *α* level set to 0.05.

#### Analysis of behavioral data

The main goal of the current study was to investigate the statistical properties of BOLD activation patterns following the presentation of fast visual object sequences. Therefore, attentive processing of all visual stimuli was a prerequisite to ensure that we would be able to decode neural representations of the stimuli from occipito-temporal fMRI data. If behavioral performance was low, we could expect that participants did not attend well to the stimuli. We thus calculated the mean behavioral accuracy on sequence and repetition trials and excluded all participants that had a mean behavioral accuracy below the 50% chance level (Fig. S1a). Mean behavioral accuracy scores of the remaining participants in the final sample are reported in the main text (Fig. 1d–f). In order to assess how well participants detected upside-down stimuli on slow trials, we conducted a one-sided one-sample t-test against the 50% chance level, testing the a-priori hypothesis that mean behavioral accuracy would be higher than chance (Fig. 1a). Cohens’d quantified the effect size and was calculated as the difference between the mean of the data and the chance level, divided by the standard deviation of the data [123]. As low performance in this task condition could be indicated by both false alarms (incorrect response to upright stimuli) and misses (missed response to upside-down stimuli) we also checked whether the frequency of false alarms and misses differed (Fig. S1b). Furthermore, we assessed if behavioral accuracy on slow trials used for classifier training was stable across task runs (Fig. S1c). In order to examine the effect of sequence speed on behavioral accuracy in sequence trials, we conducted a LME model including the sequence speed condition as the main fixed effect of interest and by-participant random intercepts and slopes. We then examined whether performance was above chance for all five speed conditions and conducted five separate one-sided one-sample t-tests testing the a-priori hypothesis that mean behavioral accuracy would be higher than a 50% chance-level. All *p* values were adjusted for multiple comparisons using the FDR-correction [129]. The effect of serial position on behavioral accuracy is reported in the SI (Fig. S1e). For repetition trials with forward and backward interference we conducted separate one-sided one-sample t-test for each repetition condition to test the a-priori hypothesis that behavioral accuracy would be higher than the 50% chance level. Results for all repetition conditions are reported in the SI (Fig. S1d). The effect sizes (Cohen’s *d*) were calculated as for slow trials.

## Data availability statement

The MRI data that support the findings of this study will be made available on https://openneuro.org/upon publication.

## Code availability statement

Custom code for all analyses conducted in this study will be made available on https://github.com/ upon publication.

## Acknowledgements

This work was funded by a research group grant awarded to NWS by the Max Planck Society (M.TN.A.BILD0004). We thank Eran Eldar, Sam Hall-McMaster and Ondřej Zíka for helpful comments on a previous version of this manuscript, Gregor Caregnato for help with participant recruitment and data collection, Anika Löwe, Sonali Beckmann and Nadine Taube for assistance with MRI data acquisition, Lion Schulz for help with behavioral data analysis, Michael Krause for support with cluster computing and all participants for their participation. LW is a pre-doctoral fellow of the International Max Planck Research School on Computational Methods in Psychiatry and Ageing Research (IMPRS COMP2PSYCH). The participating institutions are the Max Planck Institute for Human Development, Berlin, Germany, and University College London, London, UK. For more information, see https://www.mps-ucl-centre.mpg.de/en/comp2psych.

## Supplementary Information

### Additional behavioral results

Attentive processing of the visual stimuli was a prerequisite to study the evoked activation patterns in visual and ventral temporal cortex. We therefore excluded all participants that performed below chance on either or both the repetition and sequence trials of the task. To this end, we removed all participants with a mean behavioral accuracy below the 50% chance level from all further analyses (Fig. S1a). We compared the relative proportion of misses and false alarms for each of the eight functional task runs in the experiment. To this end, we conducted a LME model with trial type (miss, false alarm), session (first, second) and session run (run 1–4) as fixed effects and included by-participant random intercepts and slopes. As shown in Fig. S1b, misses (*M* = 0.55%) consistently occurred more frequently than false alarms (*M* = 0.30%), *F*_1,501.00_ = 4.1, *p* = .04, which was consistent across task runs (no effects of session or run, *p*s ≤ .70). Our classification was performed using a leave-one-run-out approach. In order to examine whether the accuracy of behavioral performance on slow trials was stable across all task runs of the study, we conducted a LME model that included the eight task runs as the fixed effect of interest as well as random intercepts and slopes for each participant. The results showed no effect of task run indicating that the accuracy of behavioral performance was relatively stable across task runs, *F*_1,92.72_ = 0.13, *p* = .72 (Fig. S1c). We examined whether behavioral accuracy on sequence trials was influenced by either the sequence speed or the serial position of the cued target image. A LME model including the sequence speed as a fixed effect and by-participant random intercepts and slopes indicated slightly lower but clear above-chance performance if the sequences were displayed at faster speeds, *F*_1,35_ = 4.27, *p* = .05 (Fig. 1f). A separate LME model including the target position as a fixed effect and by-participant random intercepts and slopes indicated lower but above-chance performance if the target image appeared at earlier serial positions, *F*_1,42.022_ = 9.92, *p* = .003 (Fig. S1d). We focused the analysis of repetition trials on the forward and backward interference condition in the main text, but also examined performance for all intermediate repetition conditions and conducted a LME model with repetition condition as a fixed effect and by-participant random intercepts and slopes. Mean behavioral performance decreased with the number of second item repetitions, *F*_1,39_ = 57.43, *p* < .001 (Fig. S1e). A series of eight one-sided one-sample t-tests indicated that for all repetition conditions mean behavioral accuracy was above the 50% chance level (*p*s ≤ .01, FDR-corrected; *d*s ≥ 0.39).

### Additional information on single event and event sequence modelling

As reported in the main text, we described multivariate decoding time courses on slow trials by a sine wave response function that was fitted to the decoding time courses of all participants separately. Evaluating a single sine wave response function for three randomly selected example participants based on the individually fitted parameters indicated that the response functions capture the individual participant data well (Fig. S2a). Based on the mean parameters across all participants we derived the mean response functions for each stimulus class which looked qualitatively similar (Fig. S2b).

**Figure S1:**
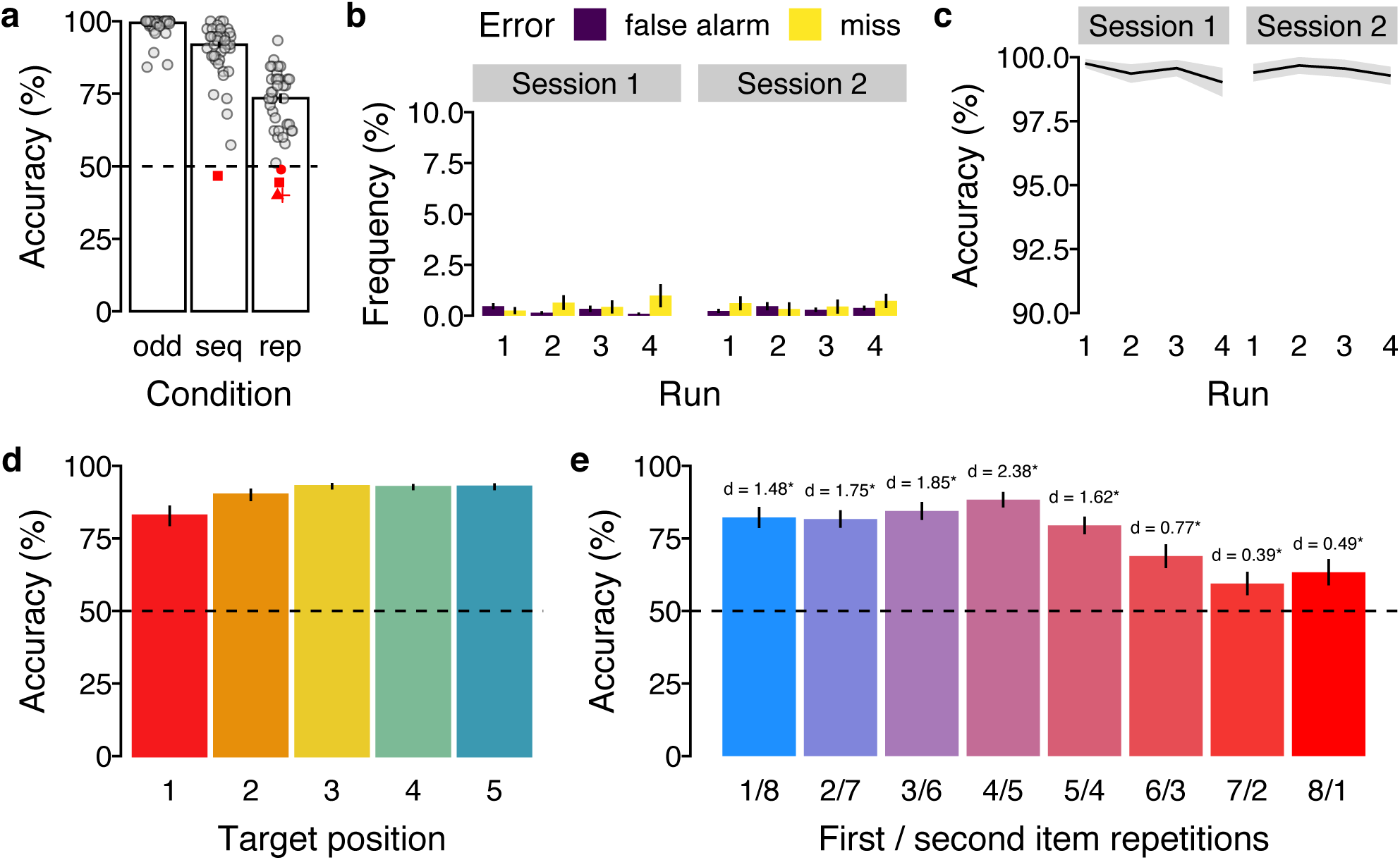
Additional behavioral results. **(a)** Mean behavioral performance (in %; y-axis) for the three trial conditions (x-axis). Dots / symbols represent mean data of one participant with below-chance performance colored in red. Note, that the SEM indicated by the errorbars was calculated after participants with below-chance performance were excluded. **(b)** Mean frequency of incorrect slow trials (in %; y-axis) across the four task runs (x-axis) of each study session (panels), separately for false alarms (violet bars) and misses (yellow bars). **(c)** Mean accuracy on slow trials (in %; y-axis) across the four task runs (x-axis) of each study session (panels). **(d)** Mean behavioral accuracy on sequence trials (in %; y-axis) as a function of serial target position (x-axis). **(e)** Mean behavioral accuracy on repetition trials (in %; y-axis) for all repetition conditions (x-axis) compared to chance. Asterisks indicate *p* < .05, FDR-corrected. Effect sizes are indicated by Cohen’s *d*. Horizontal dashed lines (in a, d, e) indicate 50% chance level. Errorbars (in a, b, d, e) and shaded areas (in c) represent ±1 SEM.

### Additional results for sequence trials

As reported in the main text, we investigated whether sequence order was evident in the relative pattern activation strength within a single measurement (i.e., within a single TR) and quantified sequential ordering by the slope of a linear regression between serial events and their classification probabilities. In addition, we repeated the same analysis using two different indices of linear association which produced qualitatively similar results. First, using ranked correlation coefficients (Kendall’s *τ*) between the serial event position and their classification probabilities as the index of linear association, we also found significant forward ordering in the forward period at sequence speeds of 128, 512 and 2048 ms (*t*s ≥ 2.22; *p*s ≤ .04, FDR-corrected; *d*s ≥ 0.37) and significant backward ordering in the backward period for all speed conditions (*t*s ≥ 4.55; *p*s ≤ .001, FDR-corrected; *d*s ≥ 0.76; Fig. S3a–b). Second, we ordered the probabilities at every TR and calculated the mean step size (i.e., difference) between the probability-ordered event positions. Again, this analysis revealed qualitatively similar results, as we found significant forward ordering in the forward period at sequence speeds of 128, 512 and 2048 ms (*t*s ≥ 2.32; *p*s ≤ .03, FDR-corrected; *d*s ≥ 0.39) and significant backward ordering in the backward period for all speed conditions (*t*s ≥ 5.17; *p*s ≤ .001, FDR-corrected; *d*s ≥ 0.86; Fig. S3c–d).

**Figure S2:**
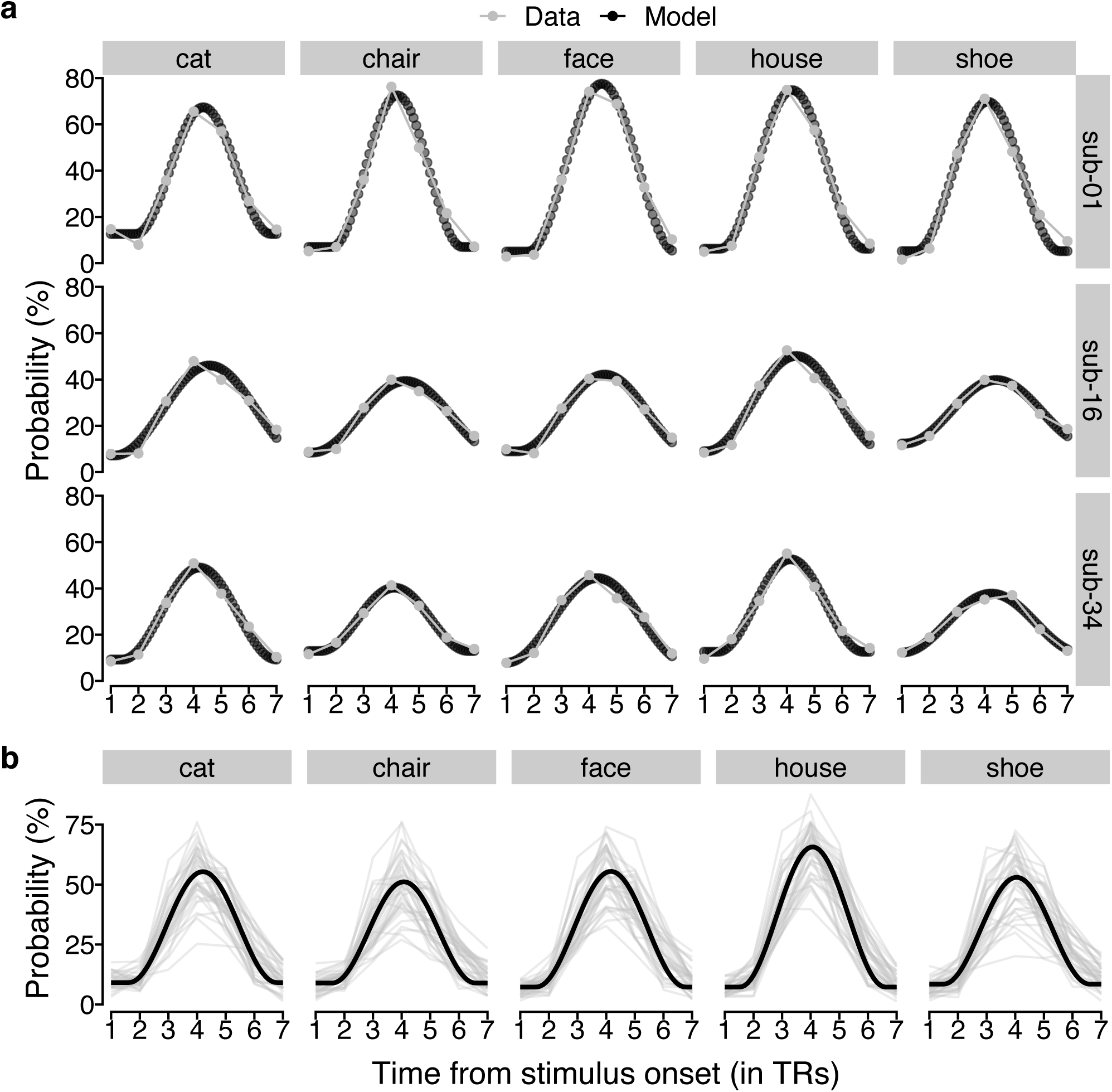
Individual fits of sine wave response function to probabilistic classifier evidence. **(a)** Time courses (in TRs from stimulus onset; x-axis) of probabilistic classifier evidence (in %; y-axis) generated by the sine wave response function with fitted parameters (black dotted line) or the true data (gray line and dots) separately for the five stimulus classes (vertical panels) and three randomly chosen example participants (horizontal panels). **(b)** Time courses (in TRs from stimulus onset; x-axis) of mean probabilistic classifier evidence (in %; y-axis) averaged separately for each participant (gray semi-transparent lines) and stimulus class (vertical panels) or predicted by the sine wave response model based on fitted parameters averaged across all participants (black line). 1 TR = 1.25 s.

Next, we analyzed the time courses of linear associations in more detail. Specifically, for each index of linear association, we tested for sequentiality at every time point (i.e., at every TR) and conducted a series of two-sided one-sample t-tests comparing the sample mean at every time point against zero (the expectation of no order information). All *p* values were adjusted for multiple comparisons by controlling the FDR across all time-points within the forward and backward period and speed conditions (38 comparisons in total). This analysis produced consistent results for each index of linear association that was tested. For the mean regression slopes, this analysis revealed significant forward sequentiality at specific earlier time points for all speed conditions (TR 3 at 32 ms, *p* = .048, *d* = 0.37; TRs 2 – 3 at 128 ms, *p*s ≤ .03, *d*s ≥ 0.38; TRs 3 – 4 at 512 ms, *p*s < .001, *d*s ≥ 0.98; TRs 3 – 7 at 2048 ms, *p*s ≤ .002, *d*s ≥ 0.60; all *p*s FDR-corrected for 38 comparisons) except the 64 ms speed condition (*p*s ≥ .08). Furthermore, we found significant backward sequentiality at specific later time points for all speed conditions (TRs 5 – 7 at 32 ms, *p*s ≤ .02, *d*s ≥ 0.43; TRs 5 – 6 at 64 ms, *p*s ≤ .01, *d*s ≥ 0.47; TRs 5 – 7 at 128 ms, *p*s ≤ .01, *d*s ≥ 0.48; TRs 6 – 7 at 512 ms, *p*s < .001, *d*s ≥ 0.98; TRs 8 – 12 at 2048 ms, *p*s < .001, *d*s ≥ 0.70; all *p*s FDR-corrected for 38 comparisons; S4a). As can be seen in Fig. S4b–d these results were qualitatively similar for all indices of linear association tested.

**Figure S3:**
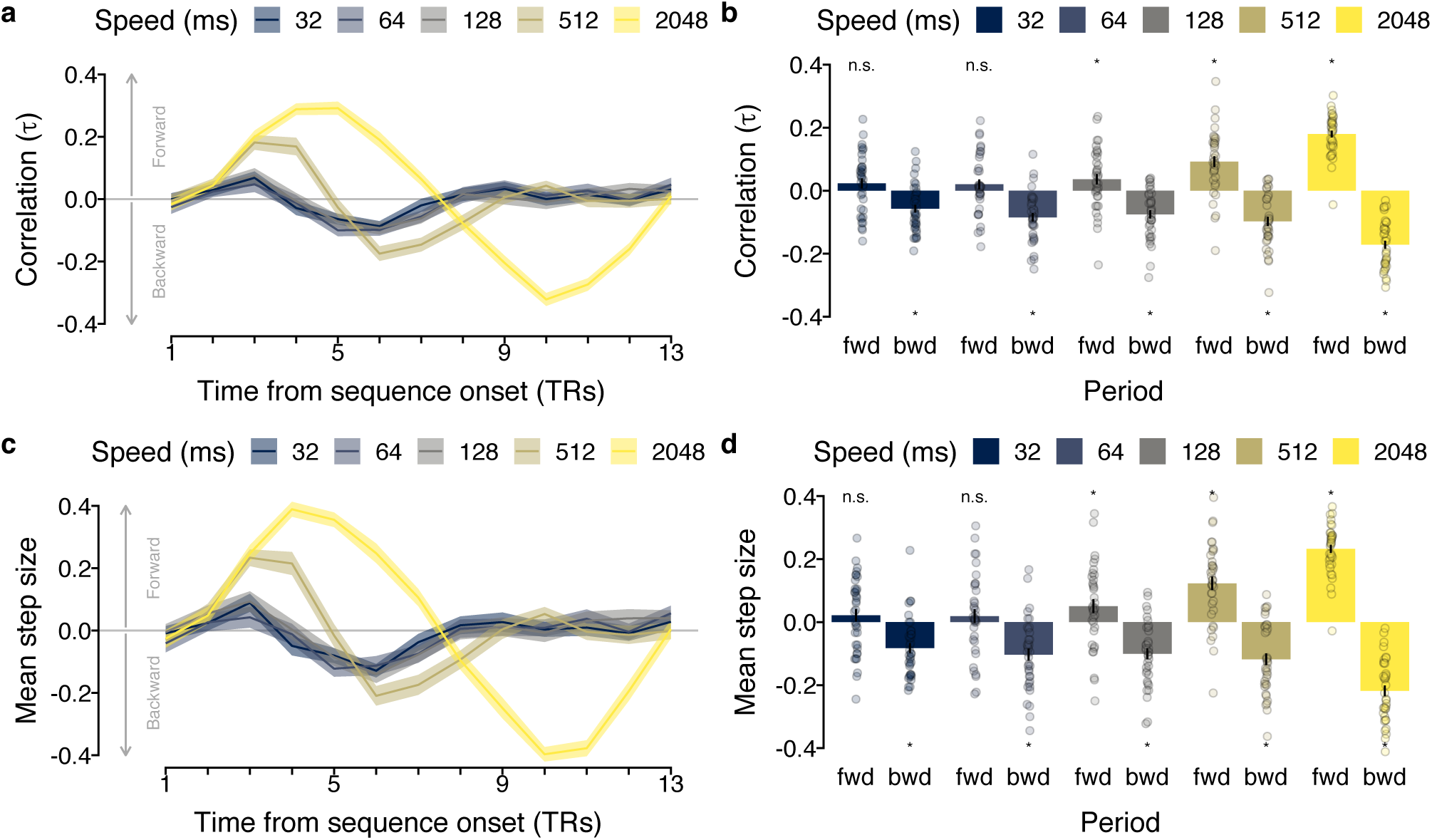
**(a)** Time courses (in TRs from sequence onset; x-axis) of mean ranked correlation coefficients between serial event position and classification probabilities (Kendall’s *τ* ; y-axis) for each speed condition (in ms; colors) on sequence trials. **(b)** Mean ranked correlation coefficients (Kendall’s *τ* ; y-axis) as a function of time period (forward versus backward; x-axis) and sequence speed (in ms; colors). **(c)** Time courses (in TRs from sequence onset; x-axis) of the mean step size between probability-ordered within-TR events (y-axis) for each speed condition (in ms; colors) on sequence trials. **(d)** Mean within-TR step-size (y-axis) as a function of time period (forward versus backward; x-axis) and sequence presentation speed (in ms; colors). Each dot in (b) and (d) represents averaged data of one participant. Shaded areas in (a), (c) and errorbars in (b), (d) represent ±1 SEM. 1 TR = 1.25 s. Stars indicate significant differences from baseline.

**Figure S4:**
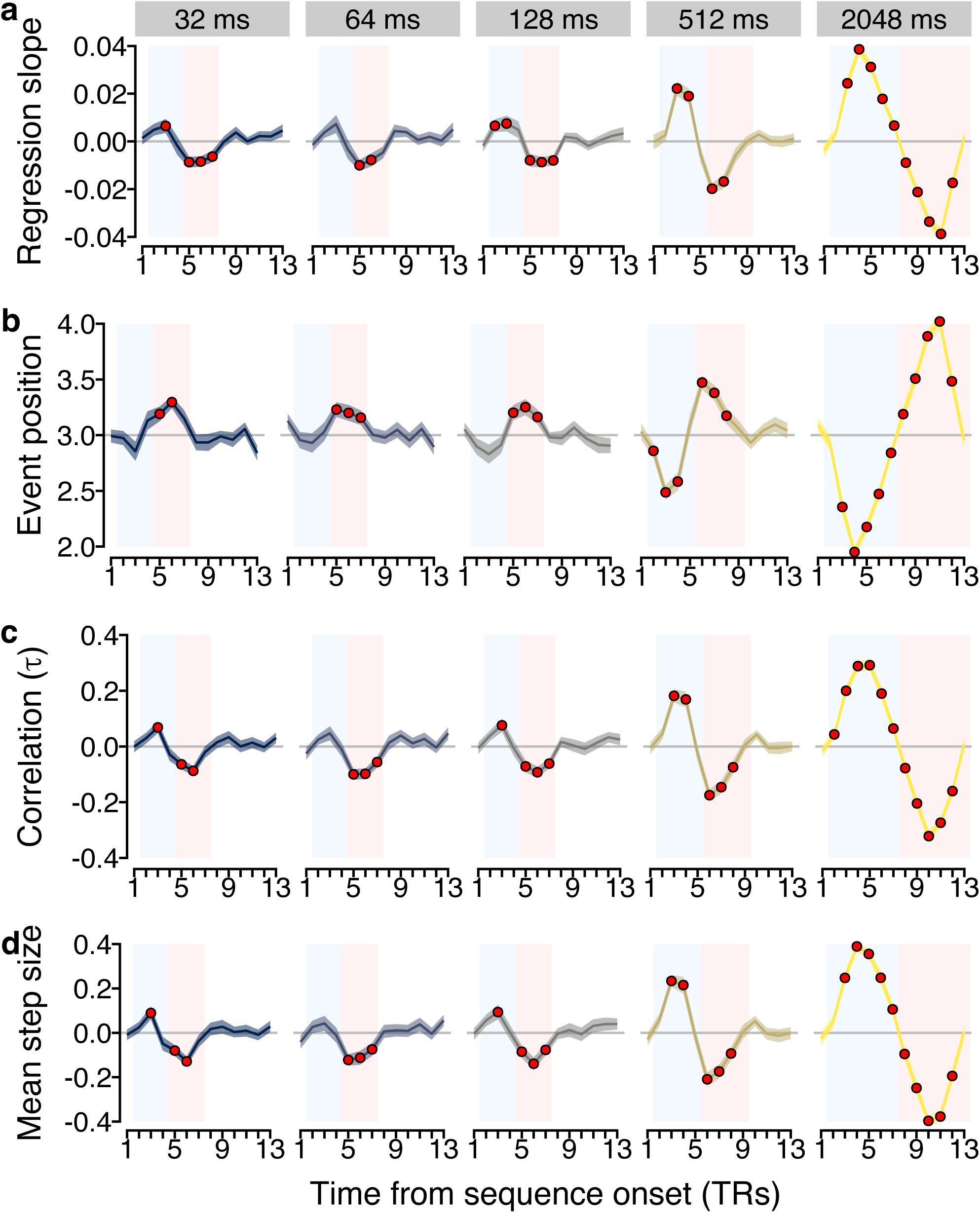
Classification time courses on sequence trials. Time courses (in TRs from sequence onset; x-axis) of **(a)** mean linear regression coefficients (slope), **(b)** mean correlation coefficients (Kendall’s *τ*), **(c)** mean step size between probability-ordered within-TR events, and **(d)** mean decoded serial event position with maximum probability for each sequence presentation speed (in ms; panels / colors). Shaded areas represent ±1 SEM. The blue and red rectangles indicate forward and backward period, respectively. Red dots indicate significant differences from baseline (horizontal gray line at zero; all *p*s ≤ .05, FDR-corrected for 38 comparisons). 1 TR = 1.25 s.

As reported in the main text, we verified that the sequentiality effects observed on sequence trials (Fig. 3b) are not only driven by the event with the maximum probability but that sequentiality is also present if the event with the maximum probability is removed. Examining the mean slope coefficients within the expected forward and backward period (adjusted by considering only four sequence events) after removing the event with the maximum probability showed that we could still find evidence for sequential ordering (Fig. S5a). Significant forward ordering in the forward period was still evident at sequence speeds of 512 and 2048 ms (*t*s ≥ 3.99; *p*s ≤ .001, FDR-corrected; *d*s ≥ 0.67) and significant backward ordering in the backward period for all speed conditions (*t*s ≥ 2.95; *p*s ≤ .009, FDR-corrected; *d*s ≥ 0.49; Fig. S5b) except the 128 ms speed condition (*p* = .10). The main analysis reported in the Results section highlighted an apparent asymmetry in detecting forward and backward sequentiality. To determine the extent to which this asymmetry was driven by the first or last item in the sequence we conducted two additional control analyses by either removing the first or last sequence item from the analysis. Removing the *first* sequence item did not change the observed sequentiality effects qualitatively (Fig. S5c) as we still found significant forward ordering in the forward period at sequence speeds of 512 and 2048 ms (*t*s ≥ 6.45; *p*s ≤ .001, FDR-corrected; *d*s ≥ 1.07) and significant backward ordering in the backward period for all speed conditions (*t*s ≥ 3.05; *p*s ≤ .006, FDR-corrected; *d*s ≥ 0.51; Fig. S5d). Removing the *last* sequence item, in contrast, made any significant sequentiality disappear for speed conditions of 128 ms or faster (*p* ≥ .12), while forward and backward sequentiality were still evident at sequence speeds of 512 ms and 2048 ms (*t*s ≥ 4.57; *p*s ≤ .001, FDR-corrected; *d*s ≥ 0.76; Fig. S5e–f).

### Additional analyses of repetition trials

We conducted two additional analyses for the data on repetition trials. First, we analyzed the effect of event duration (number of repetitions) on event probability in more detail by calculating the average event probability for each event type (*first, second*, and averaged *non-sequence*) as a function of event duration (number of repetitions). Importantly, while we focused only on the two repetition conditions with the highest degree of interference before, we now also included the data from all intermediate repetition trial types. As before, we averaged the probabilities for each serial event type but this time as a function of how often each item type was repeated in any given trial. Then, in order to test how likely we were in decoding each serial event type (first, second, non-sequence), when each item was only shown briefly once, we conducted three independent pairwise two-sample t-tests comparing the mean probabilities of all three event types with one another (correcting for multiple comparisons using Bonferroni correction).

The results reported in the main text focused on the two repetition conditions with the strongest expected effects of forward and backward interference. Additionally, we characterized the effect of event duration (number of repetitions) in more detail by analyzing the average probability of event types (first, second, non-sequence) as a function of event duration also for all intermediate repetition conditions. The results revealed a main effect of event type (first, second, non-sequence), *F*_2,282.12_ = 23.46, *p* < .001 and event duration (number of repetitions), *F*_1,71.89_ = 196.71, *p* < .001 as well as an interaction between event type and event duration, *F*_2,753.00_ = 52.46, *p* < .001 (see Fig. S6). In order to further characterize the origin of this interaction, we also conceived a reduced model that did not include the data from non-sequence events. The results of this reduced model again showed a main effect of event type (first, second), *F*_1,370.98_ = 15.32, *p* < .001 and event duration (number of repetitions), *F*_1,82.32_ = 203.32, *p* < .001 but no interaction between event type and event duration, *F*_1,502.00_ = 0.0054, *p* = .94. If only shown briefly, the second event had a mean probability (*M* = 17.11%, *SD* = 5.83%) that was higher than for the first event (*M* = 12.62%, *SD* = 5.58%), *t*_(39)_ = 2.98, *p* = .005 and the averaged non-sequence items (*M* = 7.32%, *SD* = 2.74%), *t*_(39)_ = 8.95, *p* < .001 while the average probability of the first event was also higher compared to the out-of-sequence items, *t*_(39)_ = 5.80, *p* < .001 (all *p*s were adjusted for six multiple comparisons, using the Bonferroni correction). If the event duration was prolonged (eight consecutive repetitions) the second event had a mean probability (*M* = 31.11%, *SD* = 6.87%) that was significantly different from the first event (*M* = 26.13%, *SD* = 8.28%), *t*_(39)_ = 2.70, *p* = .01 and the averaged non-sequence items (*M* = 7.49%, *SD* = 2.82%), *t*_(39)_ = 18.42, *p* < .001 while the average probability of the first event was also higher compared to the non-sequence items, *t*_(39)_ = 11.91, *p* < .001 (all *p*s were adjusted for six multiple comparisons, using the Bonferroni correction).

**Figure S5:**
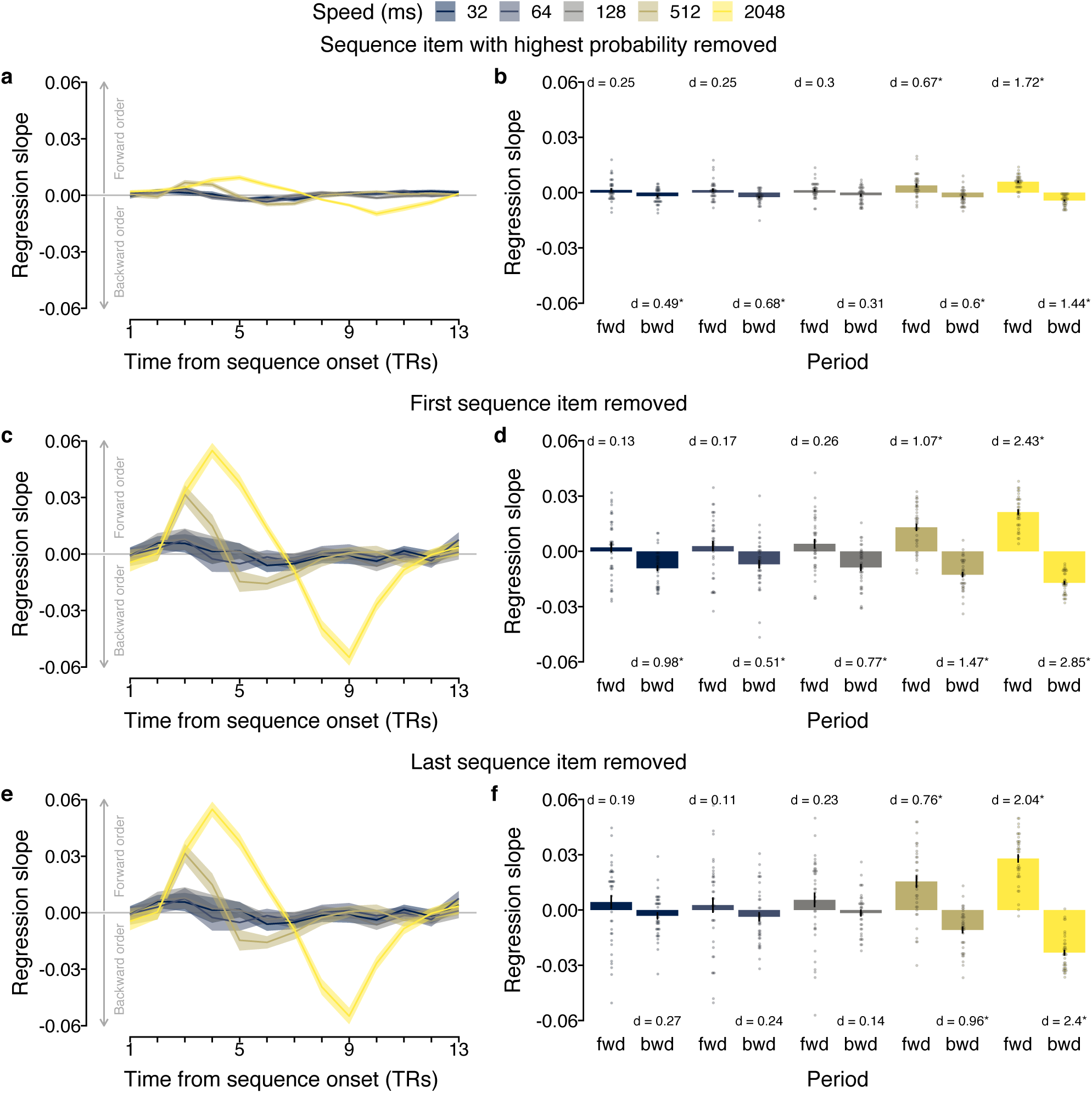
Effects of sequence item removal on sequentiality metrics. **(a, c, e)** Time courses (in TRs from sequence onset; x-axis) of mean slope coefficients of a linear regression between serial event position and classifier probability (y-axis) for each speed condition (in ms; colors) on sequence trials after removal of (a) the sequence item with the highest classification probability, (c) the first sequence item, (e) the last sequence item. **(b, d, f)** Mean slope coefficients (y-axis) as a function of time period (forward versus backward; x-axis) and sequence speed (in ms; colors) after removal of (b) the sequence item with the highest classification probability, (d) the first sequence item, (f) the last sequence item. Each dot represents averaged data of one participant. Shaded areas in (a, c, e) and errorbars in (b, d, f) represent ±1 SEM. 1 TR = 1.25 s.

These effects were attenuated but qualitatively similar when data from all TRs were considered. Specifically, a test of the model including out-of-sequence events again revealed main effects of event type, *F*_2,915_ = 14.31, *p* < .001, and event duration, *F*_1,915_ = 68.97, *p* < .001, and an interaction between the two factors, *F*_2,915_ = 17.90, *p* < .001. Testing a model without out-of-sequence events again revealed main effects of event type *F*_2,597_ = 10.92, *p* = .001, and event duration, *F*_1,597_ = 78.92, *p* < .001, but no interaction between the two factors, *F*_2,597_ = 0.18, *p* = .68. Again, the mean probability of detecting a briefly presented second (*M* = 14.41) was higher compared to a briefly presented first event (*M* = 12.02, *t*_(39)_ = 2.46, *p* = .02, Bonferroni-corrected for six comparisons). The mean probability for both briefly presented sequence items was also higher compared to out-of-sequence events (*M* = 10.28, both *t*s ≥ 2.52, both *p*s ≤ .02, Bonferroni-corrected for six comparisons). When items were repeated eight times the effect was similar: The mean probability of detecting a long second event (*M* = 19.37) was higher compared to a long first event (*M* = 16.54, *t*_(39)_ = 2.27, *p* = .03, Bonferroni-corrected for six comparisons). The mean probability for both briefly presented sequence items was also higher compared to out-of-sequence events (*M* = 9.96, both *t*s ≥ 7.99, both *p*s ≤ .001, Bonferroni-corrected for six comparisons).

**Figure S6:**
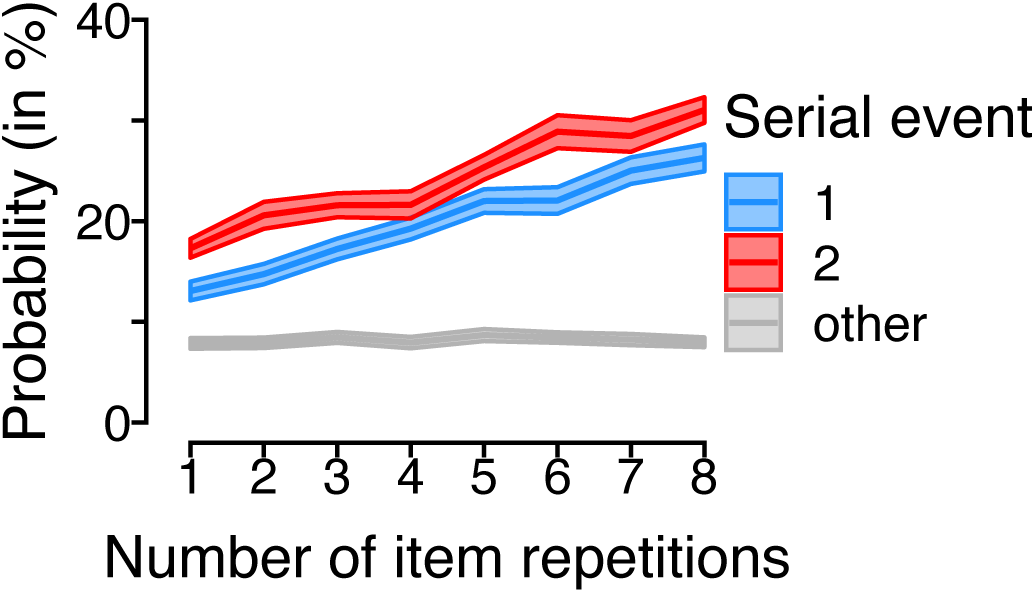
Effects of event duration (element repetition) Average probability (in %; y-axis) as a function of the number of item repetitions (i.e., total event duration), separately for event types (first, second, and out-of-sequence events; colors) based on data of all TRs.

We asked whether we would be more likely to decode items that were part of the sequence actually shown to participants (*within-sequence* items) as compared to items not part of the sequence (*out-of-sequence* items). To this end, we assessed if the serial events 1 and 2 were more likely to be decoded in the repetition trials than other events. As before, we identified the item with the highest classifier probability at every TR of each trial and then calculated the relative frequency of each item in the decoded sequence of events. These frequencies were then averaged separately for each repetition condition across all trials and participants. Next, using paired t-tests, we performed two statistical tests: First, we tested how well we were able to decode a single briefly presented item in a 32 ms sequence compared to items that were not presented, when the item is followed by a statistical representation that could mask its activation pattern (short → long trials). Second, we tested how well we were able to decode a single briefly presented item (first serial event) in a 32 ms sequence compared to items that were not part of the sequence, when the item (last serial event) is followed by a random statistical signal, for example, during an ITI (long → short trials).

Analyzing the average proportion of decoded serial events across all TRs for the *backward interference* and *forward interference* conditions separately revealed a main effect of serial event type (first, second, averaged out-of-sequence), *F*_2,234_ = 40.70, *p* = 6.80 × 10^−16^. No main effect of repetition condition (short → long versus long → short) was found, *F*_1,234_ = 0.08, *p* = .78, but an interaction between serial event position and repetition condition, *F*_2,234_ = 23.92, *p* = 3.54 × 10^−10^ (see Fig. 4e). Post-hoc comparisons indicated that in the short → long condition the longer second event had a higher frequency (*M* = 29.0%) compared to the out-of-sequence (*M* = 17.4%) as well as the short, first event (*M* = 18.9%, *ps* < .0001). The short first event did not differ from the out-of-sequence events (*p* = .47, Tukey-correction for three comparisons). In the long → short condition, in contrast, there was no difference between the long first (*M* = 24.6%) and short second event (*M* = 22.3%, *p* = .17, Tukey-correction for three comparisons) but significant differences between both within-sequence items and the averaged out-of-sequence (*M* = 17.7%) items (both *ps* < .001, Tukey-correction for three comparisons).

Analyzing the mean probability for the three event types (first, second, and out-of-sequence events) on repetition trials as a function of the absolute event occurrence per trial using data from all 13 TRs revealed a main effect of event type (first, second, out-of-sequence), *F*_2,915_ = 14.31, *p* < .001 and event duration (number of repetitions), *F*_1,915_ = 68.97, *p* < .001 as well as an interaction between event type and event duration, *F*_2,915_ = 17.90, *p* < .001 (see Fig. 4d). In order to further characterize the origin of this interaction, we also conceived a reduced model that did not include the data from out-of-sequence events. The results of this reduced model again showed a main effect of event type (first, second), *F*_1,597_ = 10.92, *p* = .001 and event duration (number of repetitions), *F*_1,597_ = 78.92, *p* < .001 but no interaction between event type and event duration, *F*_1,597_ = 0.18, *p* = 0.68. If only shown briefly, the second event had a mean probability (*M* = 14.41%, *SD* = 4.53%) that was higher than for the first event (*M* = 12.02%, *SD* = 4.78%), *t*_(39)_ = 2.46, *p* = .03 and the averaged out-of-sequence items (*M* = 10.28%, *SD* = 2.88%), *t*_(39)_ = 5.80, *p* < .001 while the average probability of the first event was also higher compared to the out-of-sequence items, *t*_(39)_ = 2.52, *p* = .03 (all *p* values were adjusted for six multiple comparisons, using the FDR correction). If the event duration was prolonged (eight consecutive repetitions) the second event had a mean probability (*M* = 19.37%, *SD* = 6.44%) that was not significantly different from the first event (*M* = 16.54%, *SD* = 4.75%), *t*_(39)_ = 2.27, *p* = .06 but from the averaged out-of-sequence items (*M* = 9.75%, *SD* = 3.05%), *t*_(39)_ = 9.36, *p* < .001 while the average probability of the first event was also higher compared to the out-of-sequence items, *t*_(39)_ = 7.99, *p* < .001 (all *p* values were adjusted for six multiple comparisons, using the FDR correction).

We also analyzed the trial-wise proportion of transition types between consecutively decoded events using data from all 13 TRs following stimulus onset. This analysis revealed that in the short → long condition the mean trial-wise proportion of forward transitions (*M* = 6.50) was higher than the mean proportion of outward transitions (*M* = 2.48), *t*_(39)_ = 4.82, *p* < .001 and also differed from the mean trial-wise proportion of outside transitions (*M* = 1.28), *t*_(39)_ = 6.14, *p* < .001 (all *p* values were corrected for four comparisons using Bonferroni correction; see Fig. 4f)). Similarly, in the long → short condition, the mean trial-wise proportion of forward transitions (*M* = 6.80) was higher than the mean proportion of outward transitions (*M* = 2.58), *t*_(39)_ = 6.11, *p* < .001 and also differ compared to the mean trial-wise proportion of outside transitions (*M* = 1.18), *t*_(39)_ = 7.71, *p* < .001 (all *p* values were corrected for four comparisons using Bonferroni correction).

#### Repeating analyses of repetition trials using data from all TRs

As reported in the main text, we focused the analyses of repetition trials on data from a relevant period of six TRs (from the second to the seventh TR) and the two trial conditions with maximum forward and backward interference, respectively. Here, we report results of the same analyses repeated using data from all TRs. The estimated probabilities of each stimulus class given the data for all repetition conditions are shown in Fig. S7. Analyzing the mean probabilities of the different event types (first, second, out-of-sequence) using data from all TRs (see Fig. S8a) revealed qualitatively similar results. Event type still influenced the average decoding probability, *F*_2,55.555_ = 41.05, *p* < .001 (see Fig. S8b). Post-hoc comparisons indicated that sequence items had a higher mean probability than out-of-sequence (9.55%) items (both *ps* < .001, Tukey-correction for three comparisons), while the second (16.77%) and first (16.77%) within-sequence event type also differed (*p* = .01, Tukey-correction for three comparisons). Repeating the analysis for the forward and backward interference conditions using data from all TRs again revealed smaller but qualitatively similar effects, with a main effect of event type (first, second, out-of-sequence), *F*_2,43.34_ = 55.42, *p* < .001, an interaction between event type and duration, *F*_2,105.00_ = 37.72, *p* < .001, and no main effect of duration (number of repetitions), *F*_1,35.70_ = 0.08, *p* = .78 (see Fig. S8c). Post-hoc comparisons indicated that in the forward interference condition the longer second event had a higher probability (19.20%) compared to both the out-of-sequence (*M* = 9.74%) and the short, first event (*M* = 11.42%, *ps* < .001, Tukey-correction for three comparisons). As reported in the main text, when using data from all TRs, the short first event did not differ from the out-of-sequence events (*p* = .13, Tukey-correction for three comparisons). In the backward interference condition, in contrast, there was no difference between the long first (16.25%) and short second event (14.34%, *p* = .22, Tukey-correction for three comparisons) but significant differences between both within-sequence items and the averaged out-of-sequence (9.36%) items (*ps* < .001, Tukey-correction for three comparisons). We also repeated the analysis investigating trial-wise proportions of transitions between consecutively decoded events using data from all TRs. Based on the full transition matrix (see Fig. S8e), this analysis revealed qualitatively similar effects (Fig. S8d): Forward transitions (3.98%) between the two sequence items were as frequent as outward transitions (2.86%, *t*_(35)_ = 2.40, *p* = .09, Bonferroni-corrected for four comparisons) but more frequent than outside transitions (2.27%, *t*_(35)_ = 3.42, *p* = .006, Bonferroni-corrected for four comparisons) in the forward interference condition. The same was true for the backward interference condition (forward transitions: 4.49%; outwards transitions: 2.89%; outside transitions: 2.35%, all *t*s ≥ 4.81, all *p*s ¡ .001; Bonferroni-corrected for four comparisons).

**Figure S7:**
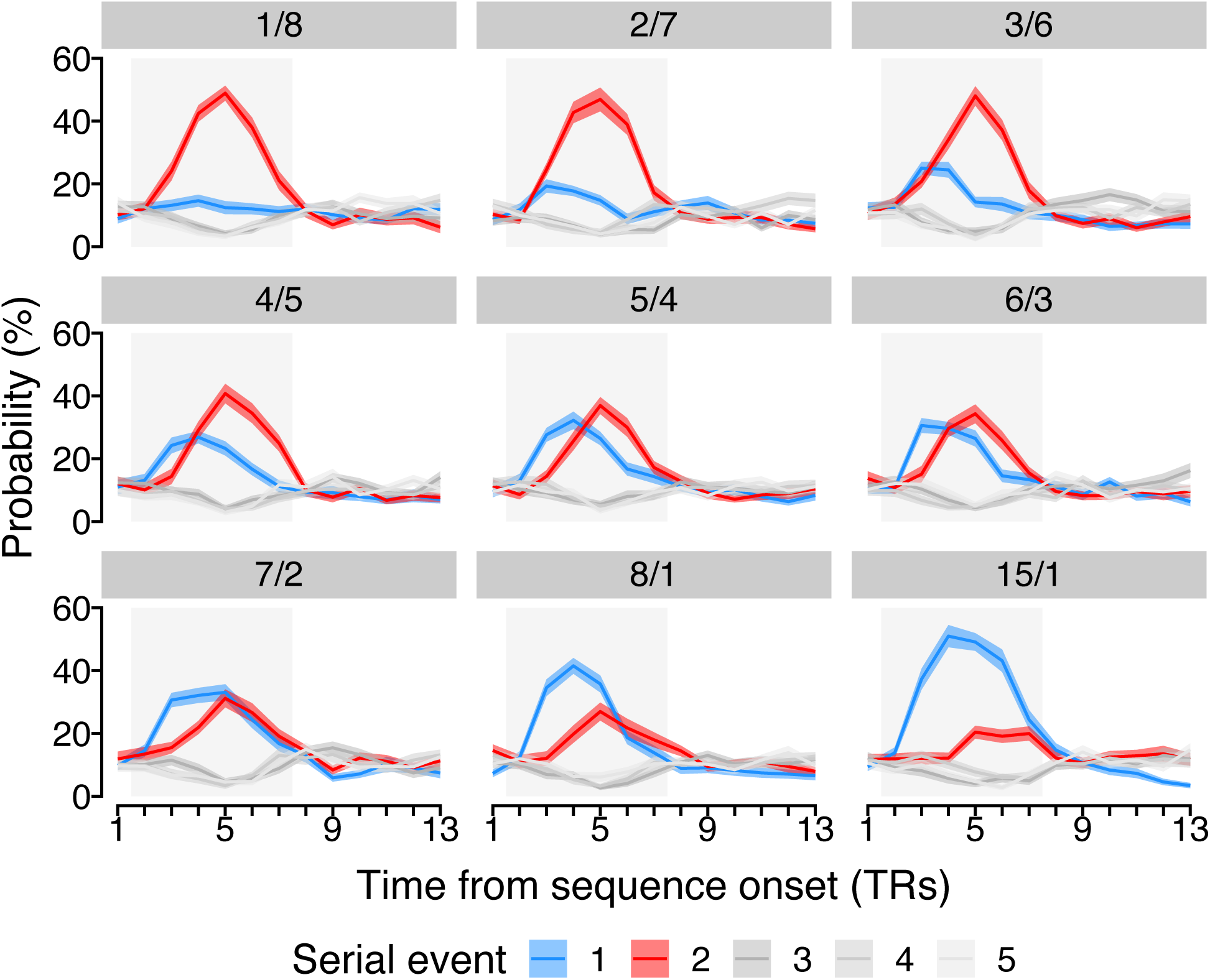
Time courses of probabilistic classifier evidence for all repetition conditions. Time courses (in TR from sequence onset; x-axis) of probabilistic classifier evidence (in %; y-axis) on repetition trials grouped by event type (colors), separately for each repetition condition (gray panels). Each panel indicates the number of repetitions per sequence event (e.g., the top-left panel indicates 1 versus 8 repeats of the first versus second event). Time-courses of classifier evidence for the first and second event are shown in blue and red, respectively, while all other stimuli that were not part of the sequence are shown in three shades of gray. Shaded areas represent ±1 SEM. 1 TR = 1.25 s.

**Figure S8:**
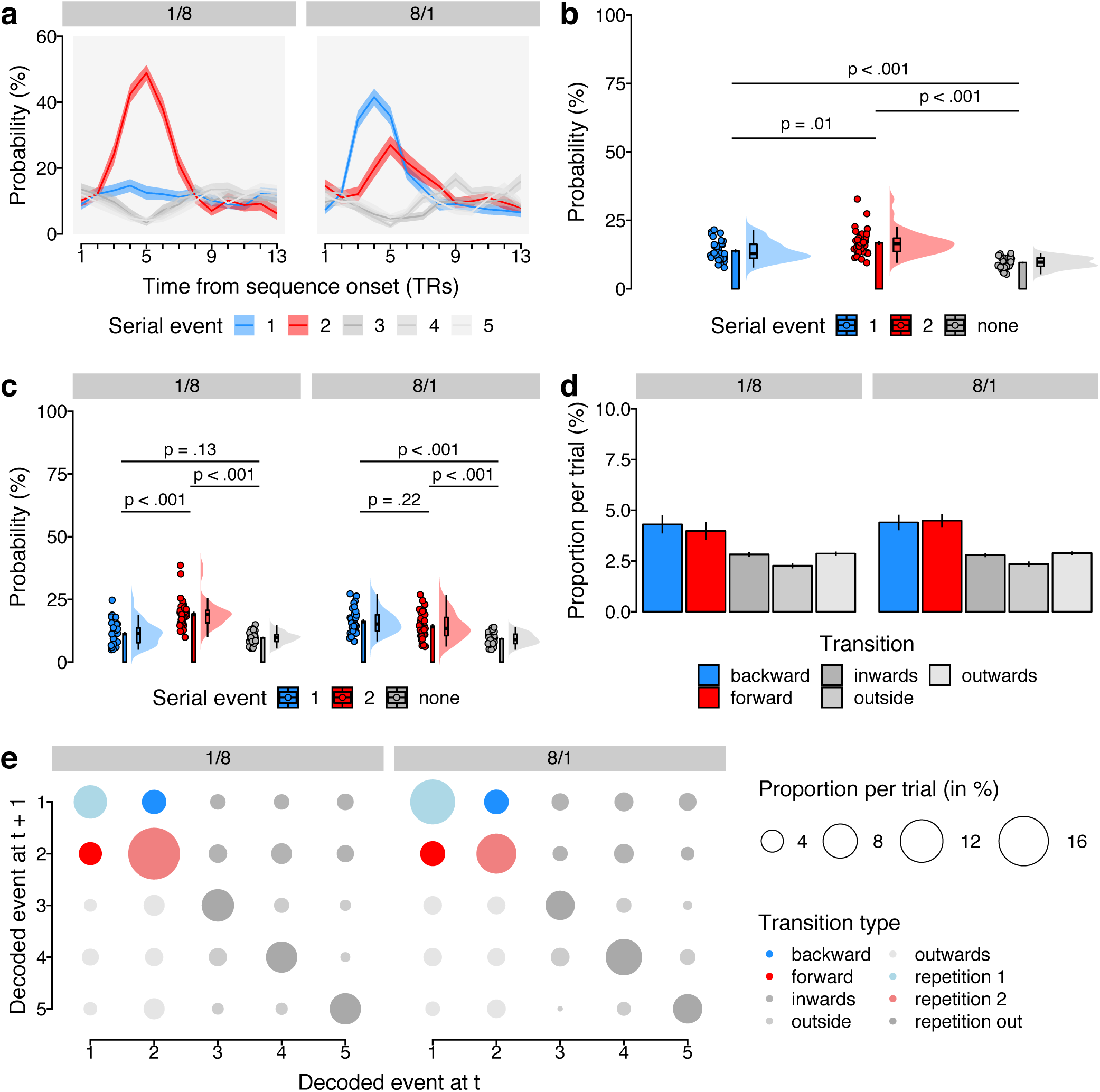
Ordering of two-item pairs on repetition trials. **(a)** Time-courses of probabilistic classifier evidence (in %; y-axis) on repetition trials as a function of time from sequence onset (in TRs; x-axis) grouped by event type (colors) for trials with backward (left panel) or forward interference (right panel). Time-courses of classifier evidence for the first and second event are shown in blue and red, respectively, while all other stimuli that were not part of the trial sequence are shown in three shades of gray. The gray rectangular area indicates the relevant time period. Ribbons represent one SEM. **(b)** Mean probability (in %; y-axis) of event types (colors) averaged across all relevant TRs). **(c)** Average probability (in %; y-axis) of event types, separately for the short → long and long → short condition (gray panels). **(d)** Mean trial-wise proportion (in %; y-axis) of each transition type, separately for the short → long and long → short condition (gray panels). **(e)** Full transition matrix of decoded event sequences indicating the mean proportion per trial (in %; circle size), separately for the short → long and long → short condition (gray panels), highlighting the transition types (colors). For all plots, each dot represents averaged data from one participant, if not indicated otherwise. The shaded areas (*rain cloud plots*) indicate the probability density function of the data [cf. 59]. The overlaid boxplots indicate the sample median alongside the interquartile range. The barplots show the sample mean and one SEM.

